# Opposing modulation of cortical and corticospinal excitability across movement-related beta stages

**DOI:** 10.64898/2026.07.28.741164

**Authors:** Marten Nuyts, Hartwig Roman Siebner, Kaat Van Dael, Lasse Christiansen, Gianmaria Senerchia, Leo Tomasevic, John Rothwell, Mikkel Malling Beck, Raf Meesen, Sybren Van Hoornweder

## Abstract

Discrete voluntary movement depends on rapid changes in neural excitability across cortical and corticospinal circuits, yet how intrinsic movement-related neural states shape multi-level excitability remains unclear. Here, we combined individualized, state-targeted transcranial magnetic stimulation (TMS) with electroencephalography and electromyography recordings during visually cued finger movements to probe excitability across movement-related beta-band dynamics during two complementary experiments. Immediate transsynaptic cortical excitability closely tracked intrinsic beta dynamics, with attenuation of the second immediate TMS-evoked potential during beta desynchronization and recovery during the post-movement beta rebound. In contrast, corticospinal excitability showed the opposite pattern, with larger motor-evoked potentials during beta desynchronization and reduced responses during beta rebound. Together, these findings identify endogenous beta-state dynamics as a key regulator of movement-related cortical excitability and reveal a fundamental dissociation between how intrinsic brain activity tunes local cortical excitability and corticospinal output in humans.

## Introduction

How the human motor system generates voluntary movement remains a fundamental question in contemporary neuroscience. Motor behavior engages local cortical circuits, large-scale networks and descending corticospinal pathways [1–3]. However, how rapid shifts in motor stages like motor preparation, execution and termination dynamically affect neural excitability remains elusive. This knowledge gap stems largely from a historical limitation: the inability to non-invasively probe multi-level human motor system excitability simultaneously and with high temporal precision.

A robust neural signature of evolving motor stages is the modulation of sensorimotor beta-band rhythms (13–30 Hz) [4, 5]. Beta oscillations are uniquely positioned to index the motor hierarchy, as they inform on sensorimotor excitability [5–7], correlate with both motor performance and clinical motor disorders [8–17], and shift predictably across the motor timeline. Specifically, beta power decreases during movement preparation and execution (movement-related beta desynchronization; MRβD) and rebounds after movement termination (post-movement rebound; PMβR) [14, 18].

Yet, relying solely on these spontaneous macro-scale read-outs leaves a critical ambiguity: beta activity alone cannot disentangle the precise, millisecond-by-millisecond excitability of the motor hierarchy. Uncovering exactly how these intrinsic, beta-defined stages shape excitability across the human motor system requires a method to actively test the system’s multilevel responsiveness. Transcranial magnetic stimulation (TMS) offers a powerful solution: a single TMS pulse over the primary motor cortex elicits an immediate local cortical response, followed by reverberant cortical and downstream corticospinal responses that collectively capture multi-level motor excitability [19–21].

At the earliest stage (< 8 ms), TMS paired with electroencephalography (EEG) uniquely captures immediate transcranial evoked potentials (iTEPs), offering a direct readout of intracortical circuit excitation [22–25]. Crucially, iTEPs are thought to reflect the cortical signature of descending corticospinal indirect (I)-waves, transsynaptic volleys generated by interneuron-mediated recruitment of corticospinal neurons [24, 26–28]. This interpretation provides a vital, non-invasive bridge to classical invasive neurophysiology findings across species [28, 29]. Subsequent TEP peaks (∼8 ms and later) reflect reverberant excitation of the motor cortex (M1) by local cortical, cortico-cortical and cortico-subcortical networks [30–32]. Finally, electromyography (EMG) captures motor-evoked potentials (MEPs), providing an aggregate measure of corticospinal excitability integrating cortical, spinal, and neuromuscular contributions [19, 33–35].

Here, we present a personalized framework that combines beta stage-targeted TMS with concurrent EEG and EMG during a visually cued finger movement task. This allows us to dissect how immediate cortical, reverberant cortical, and corticospinal excitability evolve throughout the motor timeline. Across two complementary experiments, we apply discrete stage-targeted and continuous time-resolved single-pulse TMS perturbations to directly link ongoing beta activity to immediate local cortical excitability, while simultaneously tracking how these effects propagate through the reverberant cortical network and downstream corticospinal pathways. Ultimately, this framework enables a non-invasive perturbation-based investigation into how intrinsic, oscillatory states gate multilevel excitation, revealing precisely how movement-related stage-dependent dynamics take shape across the human cortico-motor system.

## Results

To determine how intrinsic neural states shape cortical excitability across the motor timeline, we delivered suprathreshold TMS over the left primary motor hand area while concurrently recording EEG and EMG across two complementary experiments. In both experiments, healthy participants performed a visually cued, simple reaction time task consisting of a prepare cue followed by a go cue instructing a rapid right index finger keypress. In parallel, we simultaneously quantified cortical excitability across three levels: (i) immediate local cortical responses indexed by iTEPs, (ii) reverberant cortical network responses captured by TEPs, and (iii) corticospinal responses measured via MEPs.

In Experiment 1 (n = 35 recruited, n = 25 completed), we administered an individualized stimulation protocol targeting the three canonical movement-related beta stages: Rest, movement-related desynchronization (MRβD), and post-movement rebound (PMβR).

In Experiment 2, a subset of participants from Experiment 1 (n = 11 recruited, n = 10 completed) underwent a complementary time-resolved design in which neural responses were densely sampled across the entire movement cycle, together with additional resting-state measurements, to capture the continuous evolution of cortical excitability. The designs for both experiments are schematized in **Figure 1**.

**Figure 1.**
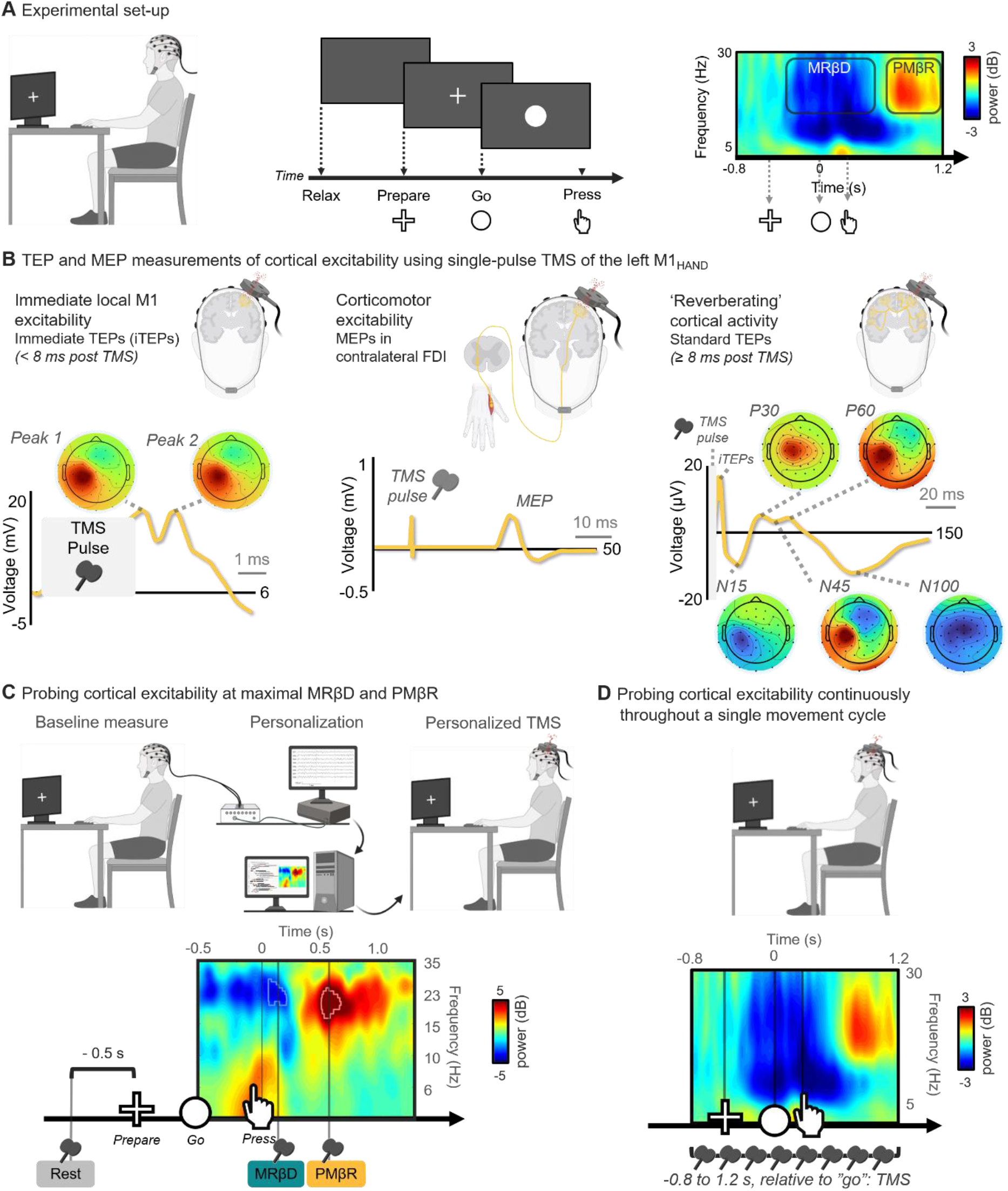
Experimental design and workflow. **A.** Experimental set-up. Participants performed a visually cued simple reaction-time task with the right index finger while electroencephalography (EEG) was used to monitor movement-related beta-band activity. Single-pulse transcranial magnetic stimulation (TMS) was applied over the hand representation of the left motor cortex (M1) to probe beta stage-dependent changes in cortical excitability. **B.** Complementary measures of cortical excitability obtained with TMS-EEG and TMS-EMG. From left to right: immediate transcranial evoked potentials (iTEPs), indexing immediate cortical excitability (< 8 ms); motor-evoked potentials (MEPs), indexing corticospinal responses recorded from the contralateral first dorsal interosseous muscle (FDI); and conventional TEPs, indexing later reverberating cortical network responses (≥ 8 ms post-TMS). EEG topographies are scaled symmetrically and depict the grand-average spatial distribution of the seven studied EEG peak responses. **C.** Experiment 1: Stage-targeted probing of cortical excitability. Subject-specific maxima of movement-related beta desynchronization (MRβD) and post-movement beta rebound (PMβR) were identified from baseline no-TMS task execution. TMS was subsequently delivered during rest, MRβD, and PMβR. **D.** Experiment 2: Time-resolved probing of cortical excitability. TMS-EEG-EMG was first acquired at rest, followed by dense sampling (-0.8 to 1.2 s relative to the go cue) throughout task execution to capture continuous stage-dependent dynamics across the movement cycle.

### Experiment 1. Canonical movement-related beta stages selectively shape cortical excitability

In Experiment 1, we used personalized TMS timings to target three canonical movement-related beta stages. While participants performed the reaction time task, single-pulse TMS was delivered at Rest, during peak MRβD and during peak PMβR, with MRβD and PMβR reflecting suppression and subsequent reinstatement of ongoing beta-band activity, respectively. Immediate cortical (iTEPs), reverberant cortical (TEPs) and corticospinal (MEPs) responses were quantified at each stage to probe beta stage-dependent changes in neural excitability (**Figure 2**).

**Figure 2.**
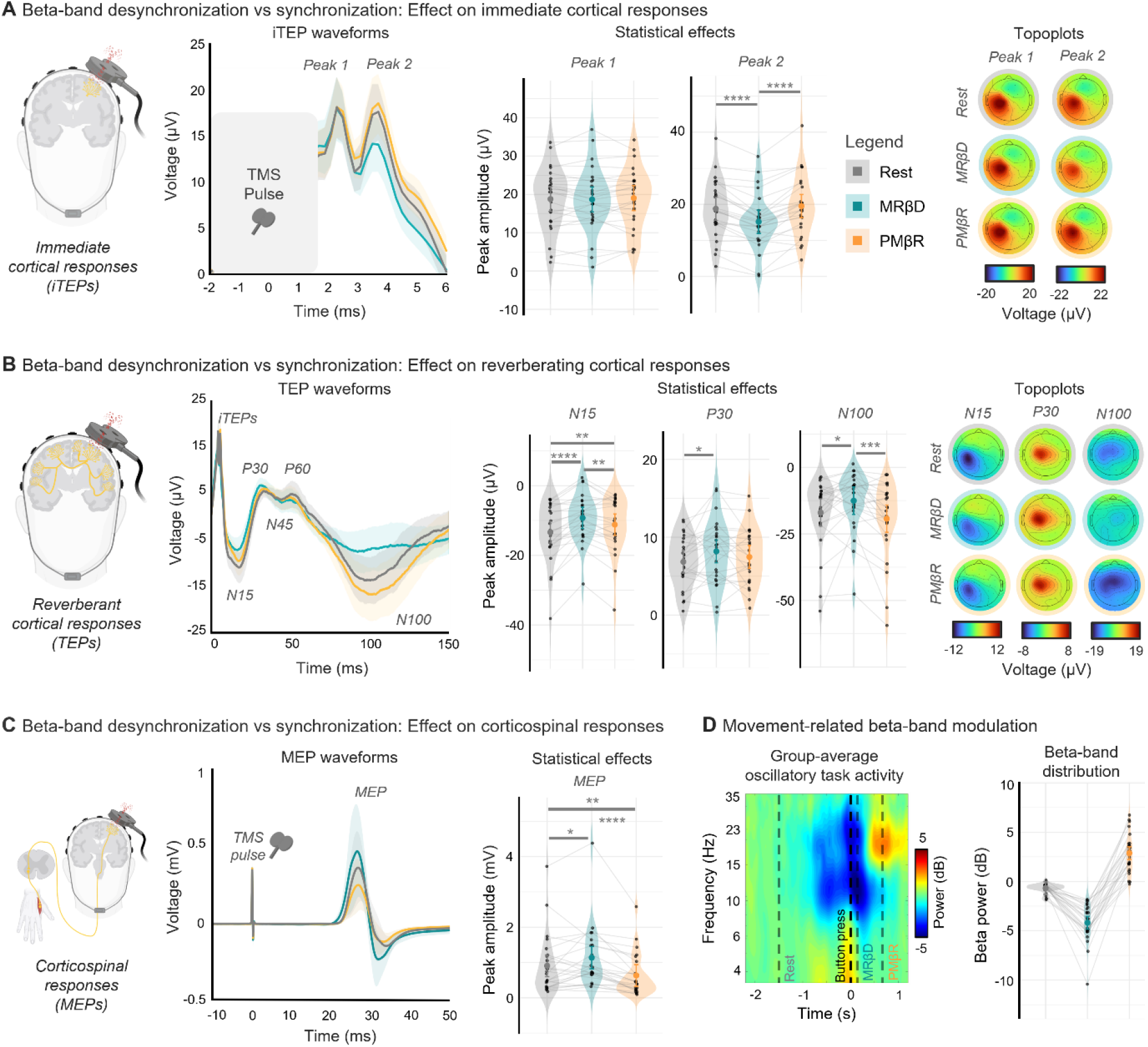
Stage-targeted transcranial magnetic stimulation (TMS) across beta-defined brain stages (Experiment 1). Panels A–C illustrate how intrinsic movement-related beta desynchronization (MRβD; blue), post-movement beta rebound (PMβR; orange), and Rest (grey) modulate responsiveness to single-pulse TMS across different levels of the motor system. For each panel, grand-average waveforms with 95% confidence intervals are shown on the left, statistical comparisons across stages in the middle, and corresponding scalp topographies on the right if applicable. **A.** Immediate cortical responses (immediate transcranial evoked potentials; iTEPs). MRβD strongly and consistently suppressed the second iTEP peak, whereas the first iTEP peak remained largely unchanged. **B.** Reverberant cortical responses (transcranial evoked potentials; TEPs). TEP responses exhibited peak-specific modulation across beta stages, most prominently for the N15 and N100 responses. **C.** Corticospinal responses (motor-evoked potentials; MEPs). Peak-to-peak MEP amplitudes were enhanced during MRβD and reduced during PMβR relative to Rest. **D.** Movement-related modulation of beta-band activity. Left: group-average time-frequency representation from an electrode over left motor cortex derived from no-TMS trials. Right: distribution of beta-band power at the time of TMS delivery, confirming successful targeting of distinct movement-related beta stages.

#### Immediate cortical responses (iTEPs)

Immediate cortical responses, thought to reflect a direct local excitation of M1 to single-pulse TMS [22, 23], were selectively modulated by ongoing movement-related beta stages (**Figure 2A**; individual traces **Figure SM1.1**). While iTEP peak 1 amplitudes remained unchanged across beta stages (F_2,48_ = 0.4, p = 0.67), iTEP peak 2 amplitudes were strongly modulated (F_2,48_ = 40.02, p < 0.001). Specifically, post-hoc testing revealed a highly consistent, significant suppression of iTEP peak 2 amplitudes during MRβD relative to Rest (*t* = -6.947, p < 0.001) and relative to PMβR (*t* = -8.355, p < 0.001). No significant differences were observed between Rest and PMβR (*t* = 1.41, p = 0.35). Together, these findings demonstrate that immediate cortical responsiveness is selectively shaped by movement-related beta stages, with iTEP peak 2 emerging as the immediate cortical response most sensitive to beta stage modulation.

#### Reverberant cortical responses (TEPs)

Reverberant cortical responses showed selective, peak-specific sensitivity to movement-related beta stages (**Figure 2B**, individual traces in **Figure SM1.2**). The early N15 peak, thought to reflect cortico-thalamo-cortical activity [36, 37], was significantly modulated across beta stages (F_2,48_ = 21.427, p < 0.001). Specifically, N15 amplitudes were significantly reduced (i.e., less negative) during MRβD compared to Rest (*t* = -6.543, p < 0.001) and PMβR (*t* = 3.081, p = 0.009), and during PMβR compared to Rest (*t* = 3.462, p = 0.003). In contrast, mid-latency P30, N45, and P60 peaks, thought to reflect reverberant excitation of M1 by local cortical, cortico-cortical and cortico-subcortical circuits [31, 32], showed limited sensitivity to beta stage. No effect was observed for N45 (F_2,48_ = 0.01, p = 0.99) or P60 (F_2,48_ = 1.97, p = 0.15), whereas a modest effect was found for P30 (F_2,48_ = 3.3, p = 0.04), driven by increased amplitudes during MRβD relative to Rest (*t* = 2.57, p = 0.035). At later latencies, beta stage yielded a strong effect on the N100 peak (F_2,48_ = 9.46, p < 0.001), a TEP response associated with large-scale integrative processes [31, 38]. N100 peak amplitudes were significantly reduced (i.e., less negative) during MRβD compared to PMβR (*t* = 4.278, p = 0.003) and Rest (*t* = 2.821, p = 0.009). Although N100 amplitudes were larger during PMβR compared to Rest, this difference did not reach significance (*t* = -1.457, p = 0.32). Together, these findings demonstrate that canonical beta stages selectively shape reverberant cortical responses, with prominent modulation of both early (N15) and late (N100) network activity.

#### Corticospinal responses (MEPs)

Corticospinal responses, reflecting the cumulative output of corticomotor projections recorded from the target muscle, were strongly modulated by beta stage (F_2,43_ = 14.72, p < 0.001) (**Figure 2C**). Peak-to-peak MEP amplitudes were enhanced during MRβD (*t* = -2.437, p = 0.049) and suppressed during PMβR (*t* = 3.228, p = 0.007) compared to Rest. Accordingly, a significant difference in MEP amplitudes was observed between MRβD and PMβR (*t* = 5.345, p < 0.001). Together, these findings demonstrate robust beta stage-dependent modulation of corticospinal responses, with facilitation during MRβD and suppression during PMβR. Notably, the corticospinal facilitation observed during MRβD co-occurred with suppression of the iTEP peak 2 and the TEP N15 and N100 peaks, indicating a surprising, differential modulation of cortical and corticospinal excitability across beta stages.

#### Coupling beta power and multilevel responses

Beyond examining discrete stages, we next tested whether individual variations in the magnitude of beta-band modulation (MRβD and PMβR relative to Rest) predicted the degree of excitability shifts across the motor system (**Figure SM1.3**). These findings reinforce the prior results, as we observed distinct, level-specific associations between oscillatory beta power shifts and cortical excitability. Individual beta-band modulation was positively coupled with iTEP peak 2 modulation: larger increases in beta power relative to Rest scaled with heightened iTEP peak 2 amplitudes (R^2^_marginal_ = 0.44, p < 0.001). Again, an opposing relationship emerged for downstream corticospinal excitability: greater beta power relative to Rest predicted a significant reduction in the amplitudes of downstream MEPs (R^2^_marginal_ = 0.13, p = 0.003). Concerning reverberant activity, greater beta power relative to rest was related to an increase (more negative voltage deflection) of the reverberant N15 (R^2^_marginal_ = 0.09, p = 0.004) and N100 (R^2^_marginal_ = 0.12, p = 0.004) TEP responses.

While these cross-stage associations hint at a continuous regulatory mechanism, confirming this in real-time requires a dense, time-resolved sampling approach across the uninterrupted movement cycle. We investigated this directly in Experiment 2.

### Experiment 2. Cortical excitability is dynamically modulated across the movement cycle

Whereas Experiment 1 established how canonical beta stage extrema shape cortical excitability, Experiment 2 introduces a complementary time-resolved framework to establish whether these relationships extend continuously throughout the movement cycle. Using a sliding-window approach, we quantified time-resolved changes in immediate cortical, reverberant cortical, and corticospinal responses from -0.8 to 1.2 s relative to the go cue during task execution (**Figure SM2.1**), together with an eyes-open resting-state baseline acquired prior to the task.

#### Immediate cortical responses (iTEPs)

Consistent with Experiment 1, immediate cortical excitability showed distinct, peak-specific modulation across the motor timeline (**Figures 3A-3C**, individual traces in **Figure SM2.2**). Most notably, iTEP peak 2 amplitude was strongly modulated, whereas iTEP peak 1 remained comparatively stable. Starting from the prepare cue, iTEP peak 2 amplitudes steadily dropped, reaching absolute minima during the go cue and the button press timing. Subsequently, there was a gradual return to pre prepare cue levels. Control analyses confirmed that these effects were not driven by underlying event-related potentials (ERPs) and were robust to baseline alignment procedures (**Figure SM2.3** and **Figure SM2.4**). To determine whether these changes reflected evolving intrinsic motor activity, we computed within-subject time-series correlations between iTEP peak amplitudes and oscillatory power across frequency bands. iTEP peak 2 showed a very strong positive relationship with beta power (mean ⍴ = 0.74, p = 0.004), whereas iTEP peak 1 showed only a moderate association (mean ⍴ = 0.41, p = 0.008) (**Figure 3D-E**). Associations with alpha- and theta-band power were substantially weaker, supporting the specificity of the relationship between immediate cortical excitability and movement-related beta dynamics. Together, these findings demonstrate that immediate cortical excitability closely tracks evolving beta dynamics, with the second iTEP peak emerging as the primary readout of beta-defined motor states.

**Figure 3.**
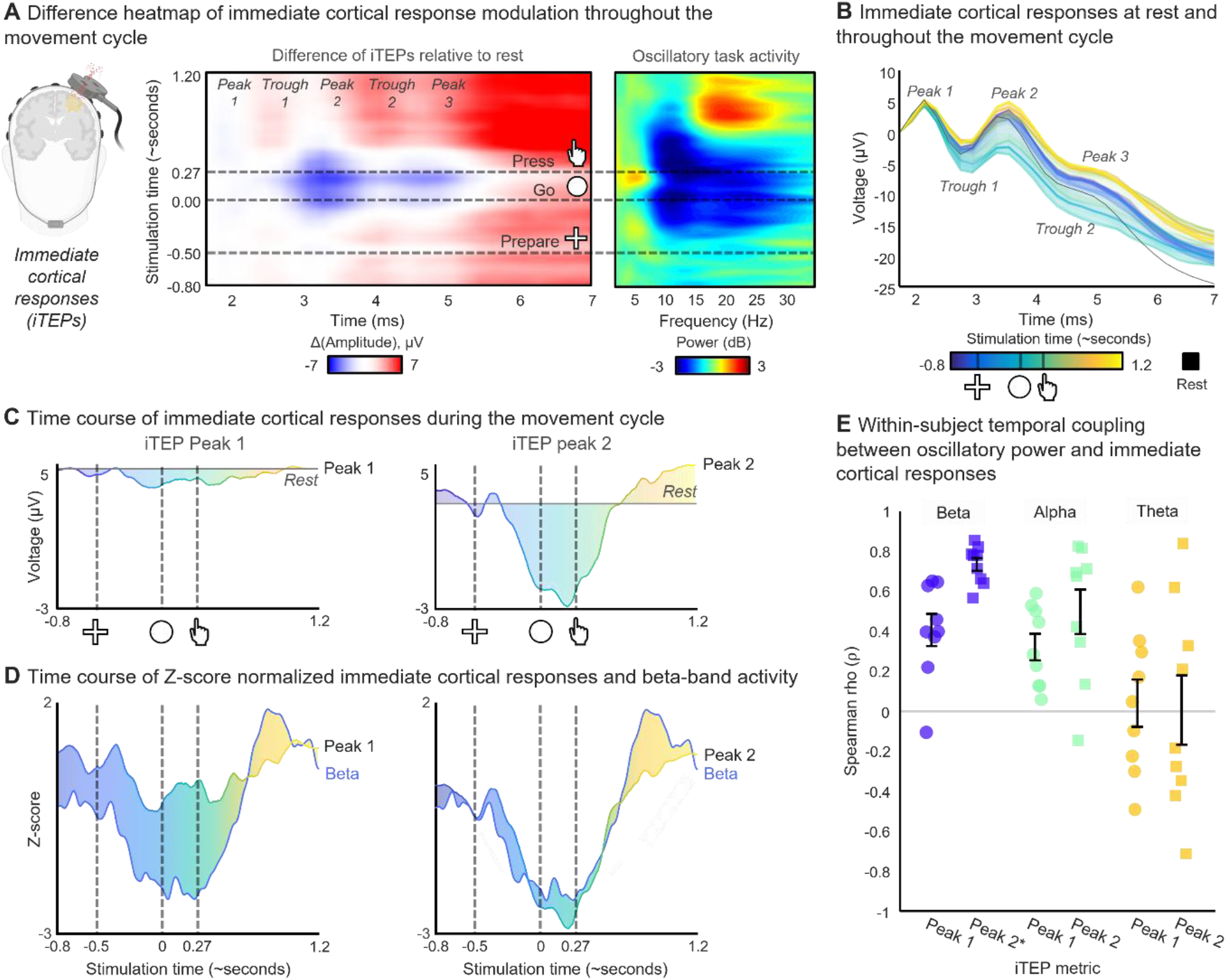
Time-resolved modulation of immediate cortical responses (Experiment 2), indexed by immediate transcranial evoked potentials (iTEPs). **A**. Difference heatmap of iTEPs relative to rest as a function of transcranial magnetic stimulation (TMS) timing (y-axis; aligned to “go” cue) and post-TMS latency (x-axis). Dashed lines indicate task events. Colors indicate increases (red) and decreases (blue) in iTEP amplitude relative to rest. Right: corresponding group-level time-frequency representation of no-TMS trials, illustrating the task-related changes in oscillatory activity. **B**. iTEP waveforms at rest (black) and across task-related TMS time points (color-coded by stimulation time). The electrode plotted is the one closest to the site of stimulation. **C**. Temporal evolution of iTEP peak magnitudes (color-coded by stimulation time) across the movement cycle. Grey line indicates resting-state baseline; dashed lines denote task events (“prepare” cue, “go” cue, button press). **D.** Z-score normalized time series illustrating the relationships between beta power (blue line) and the two iTEP peaks (color-coded by stimulation time) across the movement cycle. **E**. Within-subject correlations (Spearman ρ) between oscillatory power (beta, alpha, theta) and iTEP peak amplitude. The second iTEP peak exhibited the strongest frequency-specific coupling to beta-band activity, whereas associations with alpha and theta power were substantially weaker.

#### Reverberant cortical responses (TEPs)

Beyond the immediate cortical responses, reverberant cortical responses also exhibited distinct, peak-specific fluctuations throughout the movement cycle (**Figure 4A-C**, individual traces in **Figure SM2.5**). The N15 was consistently reduced throughout task execution relative to rest, suggesting it indexes tonic task-engagement rather than phasic motor-stage transitions. The mid-latency P30, N45, and P60 responses were broadly alike in terms of trajectories, with amplitudes increasing during movement preparation and transiently decreasing around the time of the button press. Consistent with Experiment 1, the N100 exhibited the largest modulation: amplitudes declined progressively from the prepare cue, reached a minimum just after the go cue, and subsequently increased, peaking at an absolute maximum in the period following movement termination. Control analyses confirmed that these effects were not driven by underlying ERPs and were robust to baseline alignment procedures (**Figure SM2.6** and **Figure SM2.7**).

**Figure 4.**
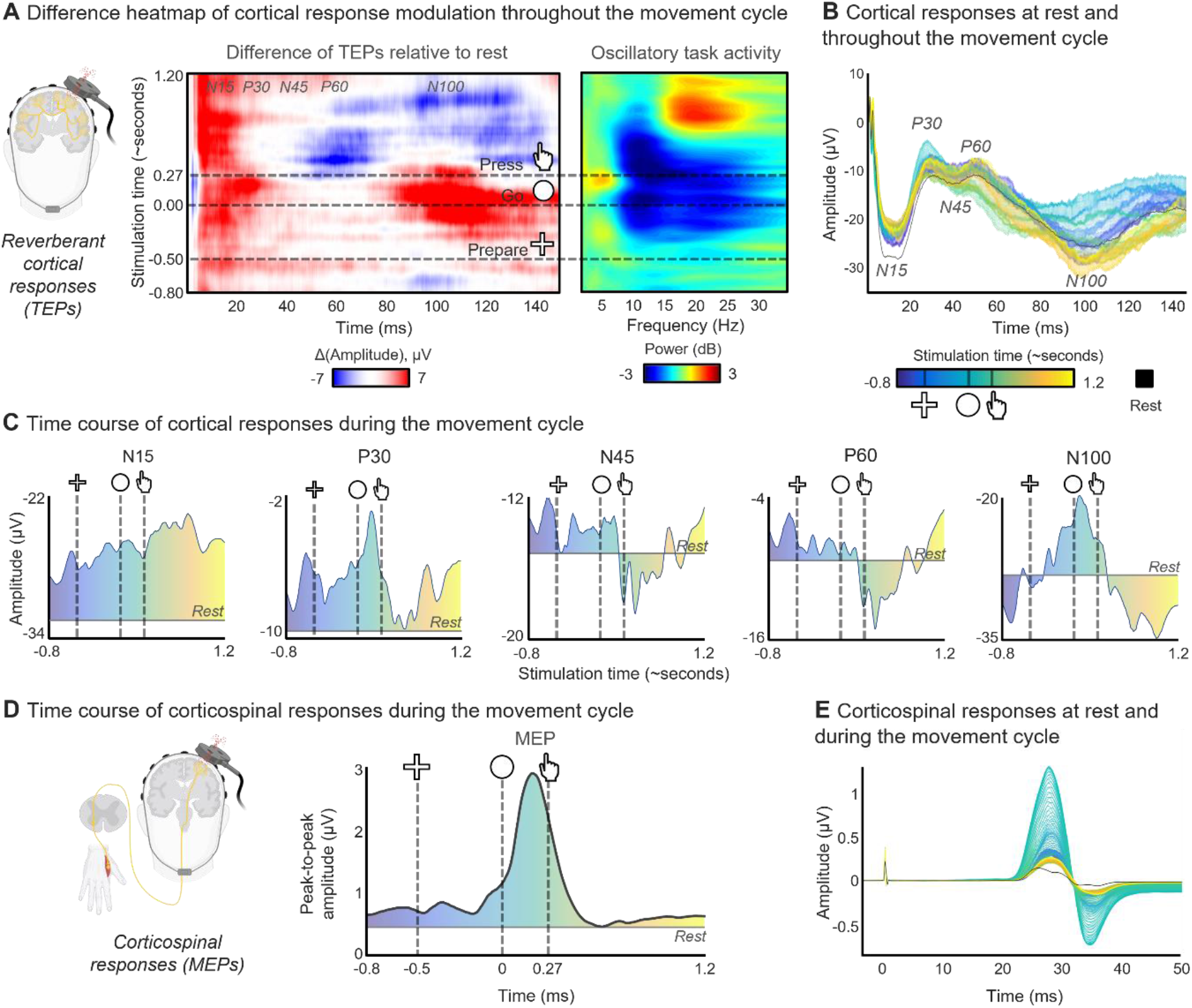
Time-resolved modulation of reverberant cortical and corticospinal responses (Experiment 2), indexed by transcranial evoked potentials (TEPs) and motor-evoked potentials (MEPs), respectively. **A.** Difference heatmap of TEPs relative to rest throughout the movement cycle (same conventions as Figure 3A). Distinct temporal modulation patterns are evident across peaks. Right: group-average time-frequency representation, illustrating the task-related changes in oscillatory activity. Dashed lines indicate task events. **B.** TEP waveforms at rest (black) and across task-related transcranial magnetic stimulation (TMS) time points (color-coded by stimulation time). The electrode plotted is the one closest to the site of stimulation. **C.** Temporal profiles of TEP peak amplitudes across the movement cycle, referenced to resting-state baseline (grey). **D.** Time-resolved modulation of MEP amplitudes, showing peak corticospinal facilitation leading up to movement excecution and suppression post-movement. **E.** MEP waveforms at rest (black) and across task-related TMS time points (color-coded by stimulation time).

#### Corticospinal responses (MEPs)

Corticospinal responses were dynamically modulated throughout the movement cycle (**Figure 4D-E**). Peak-to-peak MEP amplitudes progressively increased leading up to the button press and returned to baseline following movement termination, again showing modulation opposite to that of immediate cortical excitability. Although EMG availability was limited around movement onset (**Figure SM2.1**), the observed temporal profile concurred with established patterns of corticospinal excitability. However, we did not observe robust premovement MEP suppression, which may reflect the anticipation of some participants to the GO cue [39].

#### Coupling beta power and multilevel responses

Complementing the canonical beta stage-targeted analyses of Experiment 1, we examined whether the observed relationships between movement-related beta dynamics and cortical excitability extended continuously throughout the movement cycle. We first computed within-subject, time-resolved, Spearman correlations between beta-band power and each TMS-derived response (FDR-corrected; **Figure 5**). Subsequently, to determine how excitability evolved across the motor system, we quantified similar within-subject correlations among the immediate cortical, reverberant cortical, and corticospinal responses themselves.

**Figure 5.**
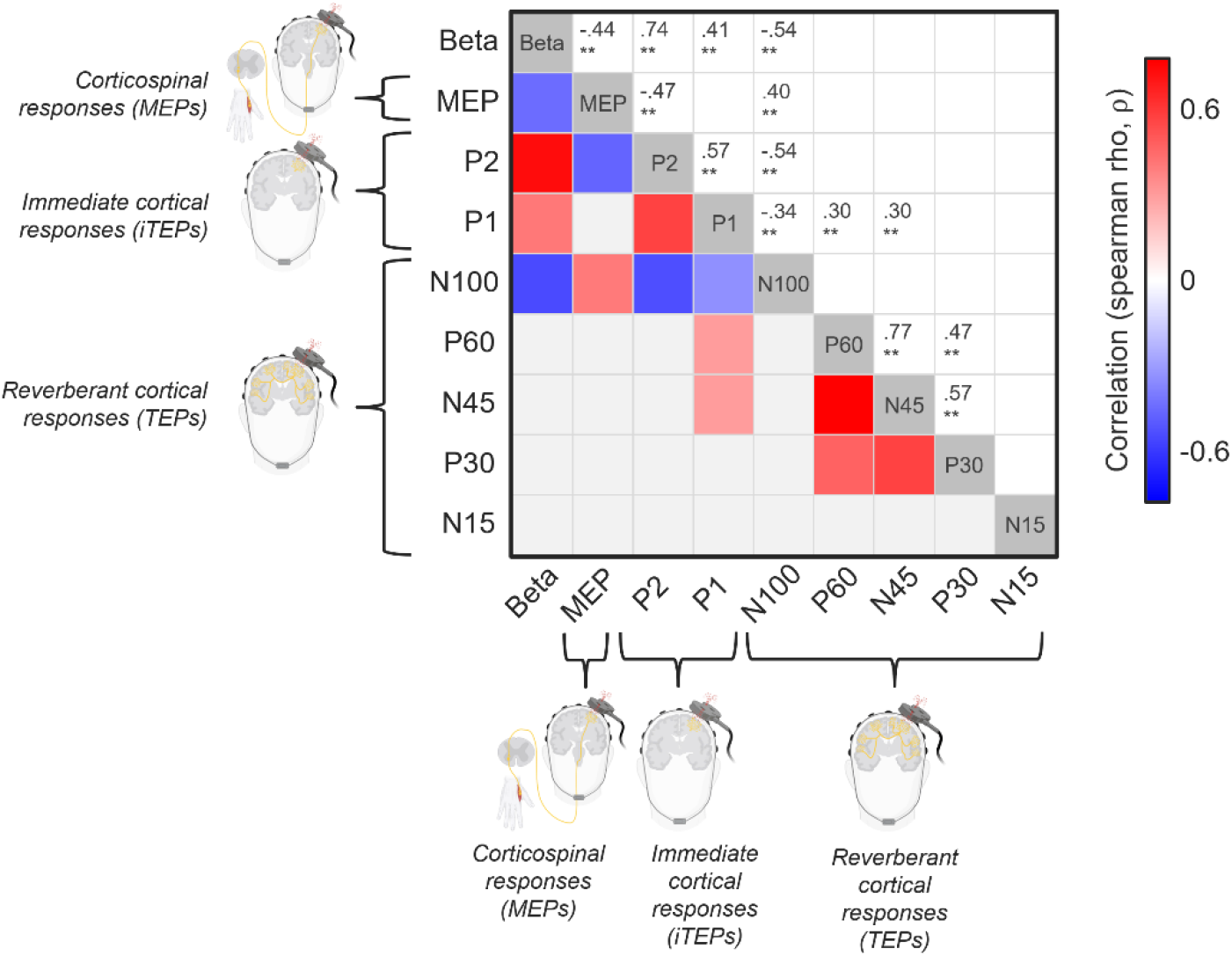
FDR-corrected correlation matrices (Spearman ρ) quantifying time-resolved coupling between (i) oscillatory beta-band power, (ii) immediate cortical responses indexed by immediate transcranial evoked potentials (iTEP) complexes, (iii) reverberant cortical responses indexed by TEP peaks, and (iv) corticospinal responses indexed by motor-evoked potentials (MEPs). A dominant cluster links beta power, iTEPs, N100, and MEPs, with the strongest association observed for iTEP peak 2. Secondary associations link between iTEP peak 2 and the N45–P60/P30 peaks, whereas the N15 remains uncoupled from other measures.

Across the motor timeline, beta power emerged as the central correlate, linking both iTEP peaks, the N100 TEP peak, and MEP amplitudes. The two iTEP peaks were positively correlated with beta power, although the association was far stronger for iTEP peak 2 (⍴ = 0.74, p = 0.004) than peak 1 (⍴ = 0.41, p = 0.008). In contrast, beta power was negatively correlated with MEP amplitudes (⍴ = -0.44, p = 0.004) and N100 amplitudes (⍴ = -0.54, p = 0.004). Notably, iTEP peak 2 was also negatively correlated with MEP amplitudes (⍴ = -0.47, p = 0.004). Together, these findings replicated the dissociation between immediate cortical and corticospinal excitability observed at the canonical beta stage extrema in Experiment 1.

A secondary cluster linked iTEP peak 1 with the N45 (⍴ = 0.30, p = 0.004) and P60 (⍴ = 0.30, p = 0.008), with peak 2 showing similar correlations to both components that did not survive FDR-thresholding. Both N45 and P60 were also strongly coupled to one another (⍴ = 0.77, p = 0.004) and correlated with P30 (⍴ = 0.57, p = 0.004, and ⍴ = 0.47, p = 0.004, respectively). The N15 was isolated from all other measures.

Together, these findings demonstrate that coupling across the motor system is non-uniform and level-specific, with immediate cortical, reverberant cortical, and corticospinal responses following partially dissociable trajectories throughout the movement cycle.

## Discussion

The present study demonstrates that movement-related beta dynamics do not exert a uniform influence across the human cortico-motor system. Instead, these dynamics differentially shape successive stages of cortical and corticospinal processing. Using a personalized TMS-EEG-EMG framework with millisecond resolution, we found that the same beta-desynchronized state that suppressed immediate cortical responsiveness simultaneously facilitated corticospinal output, a dissociation that replicated across both stage-targeted and continuous time-resolved designs. This opposing pattern is consistent with the prevailing view that movement-related beta dynamics primarily regulate the internal state and responsiveness of cortical networks [14, 18], while further demonstrating that the conversion of these cortical dynamics into descending motor output depends on additional processing across distributed cortical and subcortical circuits.

Beyond their mechanistic implications, our findings highlight the importance of multimodal approaches for studying brain-state-dependent stimulation effects. The absence of corresponding modulation of MEPs despite clear changes in early cortical responses reinforces the view that MEP amplitudes represent an aggregate physiological endpoint whose neural determinants span multiple processing stages and cannot be straightforwardly interpreted as a direct readout of cortical excitability alone [40]. Together, these findings demonstrate the value of simultaneously probing cortical and corticospinal activity when investigating the functional consequences of endogenous brain rhythms and provide an experimental framework to disentangle the distinct neural processes that collectively shape motor behavior.

A central observation is that the amplitude of the second iTEP peak closely tracked the temporal evolution of movement-related beta activity across both beta stage-targeted and continuous time-resolved experiments. This finding establishes a direct relationship between intrinsic sensorimotor beta dynamics and immediate cortical excitability. As alpha- and theta-band modulations showed substantially weaker correlations, our findings concur with prior work linking movement-related sensorimotor beta to the gating of voluntary motor output [41].

Changes in corticospinal output did not parallel the beta stage-dependent amplitude modulation of the second iTEP peak. During movement-related beta desynchronization, amplitudes of the second iTEP peak were attenuated. Conversely, MEP amplitudes increased relative to rest and post-movement beta rebound, in agreement with previous work [34, 35]. Since iTEPs are thought to reflect the cortical analogue of indirect corticospinal volleys (I-waves), which contribute to MEP generation, one would expect beta stage-dependent changes of iTEPs and MEPs to exhibit parallel dynamics. The diverging beta stage-dependent changes in iTEP peak 2 and MEP amplitude reveal a fundamental dissociation between beta-state tuning of immediate cortical excitability and corticospinal output during movement.

Several candidate mechanisms may contribute to this divergence. Movement-related beta desynchronization is generally associated with increased neuronal firing and synaptic activity within sensorimotor cortex. The concomitant amplitude reduction of the second iTEP peak may therefore reflect a state of elevated synaptic conductance in M1. By shifting local neuronal populations into a high-conductance state, increased synaptic activity could reduce the responsiveness of cortical circuits to the incoming TMS perturbation, thereby attenuating the ensuing cortical response [42]. Alternatively, beta desynchronization may be associated with greater temporal dispersion of neuronal activity within M1 microcircuits [43], reducing synchronous convergent input onto pyramidal neurons and consequently diminishing the second iTEP peak despite the concurrent increase in corticospinal excitability. Notably, attenuation of the second iTEP peak has also been reported using paired-pulse TMS paradigms probing short-interval intracortical inhibition [25], suggesting that inhibitory circuit mechanisms may contribute to this response component. Accordingly, enhanced recruitment of inhibitory M1 interneuronal networks during the beta-desynchronized state may have contributed to the observed reduction in amplitude of the second iTEP peak. Conversely, the recovery of the second iTEP peak during post-movement rebound of beta activity is consistent with the re-establishment of cortical network conditions that favor the generation of synchronized TMS-evoked responses. In parallel, the concurrent increase in MEP amplitude during beta desynchronization may reflect the combined influence of evolving cortical and spinal gain modulation [44], that are not captured by the iTEP and may partially offset or operate independently of the cortical attenuation described above.

Beyond the proposed mechanisms, the functional significance of the attenuated second iTEP peak during premovement beta desynchronization remains speculative. In this context, it is noteworthy that rapid ballistic movements are often preceded by a premovement silent period, a transient suppression of ongoing EMG activity immediately preceding the agonist burst that has been interpreted as a preparatory inhibitory process facilitating the generation of a highly synchronized and efficient motor command [45, 46]. To our knowledge, this peripheral phenomenon has not previously been linked to cortical excitability dynamics. We propose that the attenuation of the second iTEP peak during beta desynchronization may reflect an analogous preparatory process at the cortical level, transiently reconfiguring local motor-cortical networks into a state that promotes efficient and temporally coordinated corticospinal output at movement onset. If correct, this would suggest that premovement inhibitory reorganization operates in parallel at both cortical and spinal/muscular levels, as part of a broader preparatory cascade rather than two independent phenomena. This interpretation remains speculative and awaits direct empirical testing, for instance by examining whether the magnitude of iTEP 2 attenuation covaries with premovement silent period duration.

In contrast to iTEP peak 2, iTEP peak 1 remained remarkably stable across both the beta stage-targeted and continuous time-resolved experiments, exhibiting only a weak relationship with movement-related beta dynamics. This dissociation suggests that the two iTEP peaks arise from distinct physiological mechanisms. Seminal biophysical modeling work identified myelinated intracortical axons as the principal neuronal elements activated by TMS [47]. Extending this work, Worbs et al. developed a biophysical model incorporating realistic spatial patterns of intracortical and subcortical axonal myelination [48]. Using this optimized model, simulations rather suggest that axonal bends at the grey–white matter interface constitute particularly effective stimulation targets [48]. Building on these findings, we hypothesize that TMS over M1 preferentially excites cortico-cortical axons that traverse the gyral crown and bend into the superficial white matter of the precentral and postcentral gyri. Such activation would generate both orthodromic and antidromic propagation along cortico-cortical axons projecting to and from M1. Recruitment of pathways terminating within the anterior wall of the central sulcus could produce rapid activation of M1, giving rise to the first iTEP peak. Because this mode of activation may rely more strongly on direct axonal excitation than on the recruitment of local polysynaptic circuitry, it could be comparatively insensitive to fluctuations in synaptic conductance, intracortical inhibition, network synchrony, and other state-dependent processes associated with movement-related beta dynamics. This may explain the relative stability of the first iTEP peak throughout the motor timeline, in contrast to the pronounced modulation of the later transsynaptic second iTEP peak.

Although this interpretation remains speculative and will require direct testing using pharmacological interventions and circuit-specific TMS paradigms, the selective sensitivity of the second iTEP peak indicates that immediate cortical excitability is not a unitary phenomenon. Rather, distinct TMS-evoked responses appear to index different physiological processes, consistent with evidence that later I-wave-related activity is particularly sensitive to intracortical inhibitory circuitry and experimental manipulations more broadly [25, 49–51]. Reverberant cortical TEP responses at latencies exceeding 8 ms were also modulated by movement-related beta activity dynamics. The N15 and N100 TEP peaks exhibited distinct, stage-dependent modulation, indicating that these responses follow circuit-specific trajectories which are at least partially dissociable from immediate cortical and corticospinal outcomes. The early N15 TEP response was consistently attenuated during task engagement relative to the intertrial resting period, irrespective of beta stage. This finding extends the observations of Beck et al., who reported reduced N15 amplitudes during tonic muscle contraction compared with rest [36]. One possible explanation is that the early N15 response reflects TMS-evoked engagement of long-range cortico-subcortical circuits [37], whose responsiveness may be diminished in behavioral states requiring sustained corticomotor output. In contrast, the N100 response showed a clear association with movement-related beta dynamics. N100 amplitudes were consistently reduced during movement-related beta desynchronization and may indicate a transient low-gain cortical state characterized by reduced sensitivity to external input [52, 53]. A similar attenuation of the N100 response was reported by Beck et al. during tonic muscle contraction relative to rest [36]. Notably, N100 amplitudes increased after movement, paralleling the post-movement rebound in beta activity and consistent with a state of somatosensory reintegration [18, 54]. This dynamic pattern of the N100 response across movement-related beta stages is in accordance with the proposed role of N100 in large-scale integrative processing [31, 38]. Finally, the mid-latency P30, N45, P60 TEP responses exhibited comparatively weak dependence on motor stage, suggesting that these responses are less strongly linked to movement-related beta-state fluctuations than the N15 and N100 responses.

Several methodological considerations should be noted. First, although the TMS-EEG-EMG approach enables perturbational probing of human cortical physiology, mechanistic specificity remains limited relative to invasive recordings. Our non-invasive framework also precluded the assessment of beta dynamics and regional activity in subcortical structures such as the spinal cord, thalamus, subthalamic nucleus, and cerebellum which would have provided a more complete reflection of the motor system. Second, corticospinal excitability was quantified using MEP amplitudes, which are a compound measure of corticomotor projections [44]. Incorporating spinal excitability measures, such as H-reflex paradigms, would help further dissociate cortical from spinal contributions, and paired-pulse or pharmacological designs could further delineate the circuit mechanisms underlying the observed multilevel excitability modulations. Third, the current data were collected during a simple finger movement task in a restricted muscle group; whether the multilevel dissociations observed here generalize across movement types, effectors, and populations remains to be established.

In summary, we developed and applied a personalized TMS-EEG-EMG framework that links intrinsic oscillatory motor stages to multilevel cortical excitability in humans. Our complementary stage-targeted and time-resolved experiments consistently demonstrate that immediate cortical excitability closely tracks the evolution of movement-related beta dynamics, with the second iTEP peak emerging as a sensitive physiological marker of beta-defined motor stages. Beyond these immediate local cortical effects, corticospinal and reverberant cortical responses exhibited distinct patterns of stage-dependent modulation, revealing that the influence of beta dynamics is expressed differentially across levels of the motor system. Notably, corticospinal facilitation co-occurred with suppression of immediate cortical excitability during movement-related beta desynchronization, highlighting a functional dissociation between cortical input responsiveness and motor output gain. Together, these findings establish endogenous beta dynamics as a key regulator of human motor system excitability and provide causal evidence that ongoing oscillatory brain states shape neural responses to external perturbations across multiple physiological levels. More broadly, our framework provides a generalizable approach for probing state-dependent neural responsiveness in the human motor system, opening new avenues to study the pathophysiology of movement disorders and for developing neurophysiological informed brain stimulation therapies that aim to restore stage-specific neural dynamics to promote motor rehabilitation.

## Methods

### Participants

Thirty-five healthy volunteers (14 females; mean age: 22; range: 20–28) were recruited, reporting no contraindications for TMS or EEG, nor a history of neurological or psychiatric disease. All participants provided written informed consent in accordance with the Declaration of Helsinki. The study was approved by the medical ethics committee of Hasselt University (B1152024000018).

### Electroencephalography (EEG) recordings

EEG recordings were obtained from 61 passive Ag/AgCl TMS-compatible electrodes embedded in an equidistant EEG cap (M10 cap layout, BrainCap TMS, Brain Products GmbH, Germany) and connected to a TMS-compatible EEG amplifier (25 kHz sampling rate; anti-aliasing low-pass filter at 10.3 kHz; actiCHamp Plus 64 System, Brain Products GmbH, Germany). Ground and reference electrodes were placed on the left and right side of the participant’s forehead, respectively. Electrodes were prepared using abrasive gel and consistently monitored to ensure impedance levels below 5 kΩ. EEG data were visualized online and saved using BrainVision Recorder software (Brain Products GmbH, Germany). Participants were instructed to relax their face and minimize blinking throughout the experiment.

### Transcranial magnetic stimulation

To assess excitability of the left motor cortex, single biphasic TMS pulses were delivered using a ±1.5 mm thick foam-coated, 35-mm figure-of-eight coil (MC-B35 coil, MagVenture X100 with MagOption, MagVenture A/S, Farum, Denmark). The stimulator recharge delay was fixed at 500 ms following each TMS pulse. The coil was placed perpendicular to the scalp and oriented 45° relative to the midline, inducing an anteroposterior to posteroanterior current in the brain, with the coil handle pointing backwards. Stereotaxic neuronavigation (Brainsight®2, Rogue Research Inc, Montreal, Quebec, Canada) ensured stable and accurate coil positioning. Participants wore modified earplugs and pink noise mixed with pre-recorded TMS clicks was played via the TAAC software during data acquisition to attenuate TMS-click sound effects on the EEG recordings [55].

### Electromyography (EMG) recordings

Corticospinal responses, MEPs, were recorded via self-adhesive surface EMG electrodes (Ambu Neuroline 700), attached to the right first dorsal interosseus muscle belly and tendon, with a ground electrode placed over the right ulnar styloid process. The EMG signal was amplified (gain = 1000) (Digitimer D360, Digitimer Ltd., Hertfordshire, UK), 0.02-2 kHz bandpass filtered, 5 kHz digitized (CED 1401 micro, CED Limited, Cambridge, UK), and stored for offline analyses.

### Simple reaction time task

Using Psychtoolbox-3 and MATLAB (2022b, The Math-Works Inc., Portola Valley, California, USA), a visually cued simple reaction time task was employed [56]. Participants were seated comfortably and instructed to place their right index finger on the space bar of a keyboard. Each trial consisted of three stages: rest, preparation and execution. During the preparation stage, a fixation cross was presented on a screen in front of the participant for a variable duration of 600-800 ms (Experiment 1) or 400-600 ms (Experiment 2). Next, a visual cue (circle) appeared on the screen, informing participants to press the spacebar as quickly as possible. Keypress timing was detected online using Psychtoolbox functions, with the keypress event used as the primary temporal reference for all movement-relative analyses. Following each response, a blank screen was shown for 3 s. It should be noted that USB keyboard acquisition introduces variable polling-dependent latency (typically about 8–30 ms).

### Protocol Experiment 1

An overview of the experimental protocol is shown in **Figure 1B**. Following a short task familiarization, EEG data were recorded while participants completed 60 trials of the simple reaction time task (i.e., baseline measurement). These data were automatically processed offline to derive each participant’s peak MRβD and PMβR timing following the button press (see Supplementary Materials 3 for pipeline details). These individual timings were later used to personalize TMS timings (**Table SM3.1**).

In parallel, several TMS procedures were performed. The motor hotspot of the right first dorsal interosseous (FDI) muscle was determined by moving the coil in small steps to identify the spot with the largest and most consistent MEPs. Over the hotspot, the resting motor threshold (rMT) was defined as the minimal intensity required to evoke MEPs with a peak-to-peak amplitude of ≥ 50 µV in at least five out of ten consecutive trials. To enable iTEP detection, we followed a previously established hotspotting procedure [22]. Briefly, 10 to 20 suprathreshold pulses (110% rMT) were delivered over the motor hotspot and the resulting averaged EEG response was inspected online using BrainVision Recorder [57]. When scalp muscle artifacts were present, which was the case in 17 participants, small adjustments in coil tilt and/or position were made until these artifacts were absent while MEPs remained present. The resting motor threshold was reassessed after any adjustment.

In twenty-five participants (11 females; mean age: 23; range: 20–28), TMS-EEG data were recorded at 110% rMT. The remaining 10 participants were excluded due to: scalp muscle co-activation despite tailored hotspotting (n = 8); participant fatigue (n = 1); or absence of MRβD during the baseline task (n = 1). TMS-EEG data were collected across five blocks, consisting of 80 trials each, while the participant performed the simple reaction time task. Trials consisted of either (i) single-pulse TMS 500 ms prior to the task onset (Rest), (ii) single-pulse TMS during an individual’s peak MRβD, (iii) single-pulse TMS during an individual’s peak PMβR, or (iv) no-TMS (Catch). A ± 50 ms jitter was added to each TMS pulse. Each condition randomly occurred twenty times per block, totaling 100 trials per condition. TMS pulses were delivered at 110% rMT. The catch trials (no-TMS) were used to extract oscillatory activity during the behavioral task. **Figure SM3.1** demonstrates that we successfully induced and targeted MRβD and PMβR.

### Protocol Experiment 2

An overview of the experimental protocol is shown in **Figure 1C**. Eleven participants (5 females; mean age: 24; range: 22-27) from Experiment 1 were reinvited for a second follow-up experiment. One participant was excluded due to technical issues. In one of the remaining participants (n =10), EEG data were of insufficient quality due to decay artifacts and/or excessive noise affecting a substantial number of channels of interest, and were therefore excluded, while EMG data were retained for analysis.

First, TMS-EEG data were recorded at 105% rMT while participants were in resting-state (eyes open), staring at a cross in front of them. Next, a short task familiarization of the simple reaction time task was performed. Subsequently, participants performed six blocks of the simple reaction time task. Each block consisted of 85 trials, of which 72 trials contained TMS and 13 were catch trials (no-TMS). The catch trials (n = 78) were used to extract oscillatory activity during the behavioral task. In the TMS trials (n = 432), pulses were applied at linearly spaced intervals between -0.8 to 1.2 seconds with respect to the “go” signal. Overall, this approach was successful in allowing for time-resolved sampling of the movement cycle, as shown in **Figure SM2.1.**

### Data preprocessing

All data were preprocessed and analyzed using custom MATLAB scripts with EEGLAB [58] and Fieldtrip functions [59].

#### Experiment 1

EEG data acquired during baseline and catch trials (i.e., no-TMS data) were downsampled from 25 to 1 kHz, band-pass filtered (1-35 Hz), and re-referenced to the common average reference. Independent component analysis was performed to remove ocular and/or muscular activity, identified with a probability exceeding 90% [60]. EEG data from sensor C3, over the left motor cortex, were decomposed into a time-frequency representation using complex Morlet wavelet convolution (4-35 Hz, in 1 Hz steps). Wavelet widths increased logarithmically from 3 to 10 cycles across frequencies [61]. Power was baseline-normalized, using a -2 to -1.5 s window relative to the button press, and expressed in decibels. Beta-band power was extracted during peak MRβD, peak PMβR, and Rest (cf., **Figure SM3.1**).

EEG data acquired during TMS trials (i.e., TMS-EEG data) were epoched from -1 to 1 s around the TMS pulse. The TMS pulse artifact was removed by cubic interpolation from -1.9 ms before to 2 ms after the TMS pulse. The data were then band-pass filtered (2nd order zero-phase Butterworth filter at 0.1-2,000 Hz) and downsampled from 25 to 5 kHz. Individual trials were visually inspected for ocular and/or muscular activity, and contaminated trials were discarded (median: 8; range: 0-35). Channels containing excessive noise (median: 1; range: 0-6) and/or decay artifacts (median: 1; range: 0-3) were interpolated. Finally, the data were baseline-corrected from a window of -110 ms to -10 ms relative to the TMS pulse onset. In Experiment 1, canonical (i)TEP peaks were extracted per beta stage (Rest, MRβD, and PMβR) from the averaged signal (see Supplementary Materials 4 for peak extraction details), from the electrode closest to the site of stimulation (**Figure SM1.4**), in line with prior work [22, 62].

For the EMG data, individual trials were discarded when the root-mean-square amplitude in the 50 ms window preceding the TMS pulse exceeded 30 µV (median: 10; range: 0-98). By doing so, no residual pre-stimulus EMG activity remained (**Figure SM3.2**). Six participants had fewer than 25 remaining trials in the MRβD condition and were therefore excluded from further EMG analyses, as such low trial numbers yield unreliable MEP estimates [63]. Peak-to-peak MEP amplitudes were calculated from the averaged EMG trace as the difference between the minimum and maximum values in the 20–40 ms window post-TMS.

#### Experiment 2

EEG data acquired during catch trials (no-TMS) were downsampled to 500 Hz, band-pass filtered (1–45 Hz), and re-referenced to the common average reference. Akin to Experiment 1, independent component analysis was performed to remove ocular and/or muscular activity and EEG data from sensor C3 were decomposed into a time-frequency representation using complex Morlet wavelet convolution (1-40 Hz, in 34 steps). Wavelet widths increased logarithmically from 3 to 10 cycles across frequencies [61]. Power was baseline-normalized, using a -1.5 to -1 s window relative to the button press, and expressed in decibels. The mean theta (4–8 Hz), alpha (8–12.5 Hz) and beta-band (13.5–30 Hz) trajectories were extracted per participant from an average of 64.22 ± 7.94 trials per participants (range: 49–73). EEG data acquired during TMS (i.e., TMS-EEG data) were processed in identical fashion as Experiment 1. Channels containing excessive noise (median: 2; range: 0-4) and/or decay artifacts (median: 2; range: 1-4) were interpolated. For the resting-state data, 55.44 ± 3.97 (range: 49–60) were analyzed. For the task-related data, 391.22 ± 25.35 (range: 326–413) were analyzed. **Figure SM2.1** shows the empirical cumulative distribution function of the trial counts.

For the EMG data, MEPs were manually preprocessed and trials where the TMS-pulse and MEP fully overlapped with ongoing muscle activity of the reaction time task were removed as no MEP could be identified. All other trials were included, resulting in 60 trials across all participants for the resting-state condition, and 313.90 ± 37.57 (range: 224–356) trials for the task condition. **Figure SM2.1** shows the empirical cumulative distribution function of the trial counts.

Trials with reaction times <150 ms were discarded, as these were considered anticipatory responses.

### Analyses

Statistical analyses were conducted in Matlab and R(Studio), with alpha set at 0.05 [64, 65]. Assumptions for each statistical test were verified prior to analysis.

#### Experiment 1

To assess beta stage-dependent modulation of cortical excitability, separate linear mixed-effects models were fitted for each TMS-evoked response, including (i) iTEP amplitudes (Peak 1, Peak 2), (ii) TEP amplitudes (N15, P30, N45, P60, N100), and (iii) MEP peak-to-peak amplitudes:

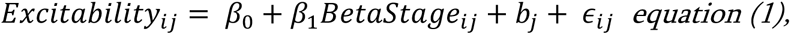

Where BetaStage (Rest, MRβD, PMβR) was modeled as a fixed effect and participant (b_j_) as a random intercept. When relevant, Tukey-corrected post-hoc comparisons were performed.

Furthermore, to assess if the degree of beta modulation with respect to rest related to any of the cortical excitability measures, we tested the following linear mixed model:

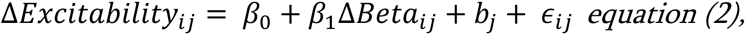

Where ΔExcitability refers to the change in excitability during MRβD or PMβR relative to Rest, and ΔBeta refers to the change in beta-band oscillatory power during MRβD or PMβR relative to Rest. Participant was included as a random intercept (b_j_) to account for the two included beta stages.

#### Experiment 2

To analyze excitability changes across the movement cycle, TMS-evoked responses were first temporally realigned to a normalized group-level movement axis using subject-specific interpolation between trial start, movement onset, and response. For each subject, trials were then sorted according to TMS timing within this normalized axis. This procedure ensured that responses were analyzed relative to task events (“prepare”, “go”, “press”), rather than absolute stimulation time, which would otherwise obscure the behaviorally relevant temporal structure.

Time-resolved excitability trajectories were computed using a sliding-window approach (window size: 30 trials, step size: 1 trial) with Gaussian weighting, yielding continuous estimates of (i) iTEP, (ii) TEP, and (iii) MEP amplitudes. These trajectories were interpolated onto a common normalized time grid (−0.8 to 1.2 s) to enable group-level comparison. Non-interpolated data are provided in Supplementary Materials 2 (**Figures SM2.8** and **SM2.9**) and highlight that our findings are robust to the choice of interpolation. For visualization, (i)TEP trajectories in the main manuscript were baseline-shifted such that the first analyzed time point (1.6 ms) was set to zero to account for minor offsets in the data; corresponding non-shifted data are provided in Supplementary Materials 2 (**Figures SM2.3**-**4** and **SM2.6-7**) and yielded identical results. In addition, control analyses correcting for the ongoing event-related potential are reported in Supplementary Materials 2 (**Figures SM2.3**-**4** and **SM2.6-7**) and did not affect the study results.

The trajectories were quantitatively assessed across the movement cycle and with respect to resting-state measures. To assess coordinated dynamics across measures, within-subject associations between time-resolved trajectories (oscillatory power and multilevel TMS-evoked responses) were computed using Spearman correlation. Prior to analysis, variables were Fisher z-transformed. Where applicable, multiple comparisons were controlled using false discovery rate (FDR) correction (**Figure 3D** and **Figure 5**).

## Data availability

The data supporting the findings of this study are available from the corresponding authors, Marten Nuyts and Sybren Van Hoornweder, upon reasonably motivated request.

