## Supplementary Materials for "Opposing modulation of cortical and corticospinal excitability across movement-related beta stages"

[1] Shared authorship

### Supplementary Materials 1. Additional results of Experiment 1.

The first part of the Supplementary Materials provides several additional analyses and visualizations that support the main results section of Experiment 1.

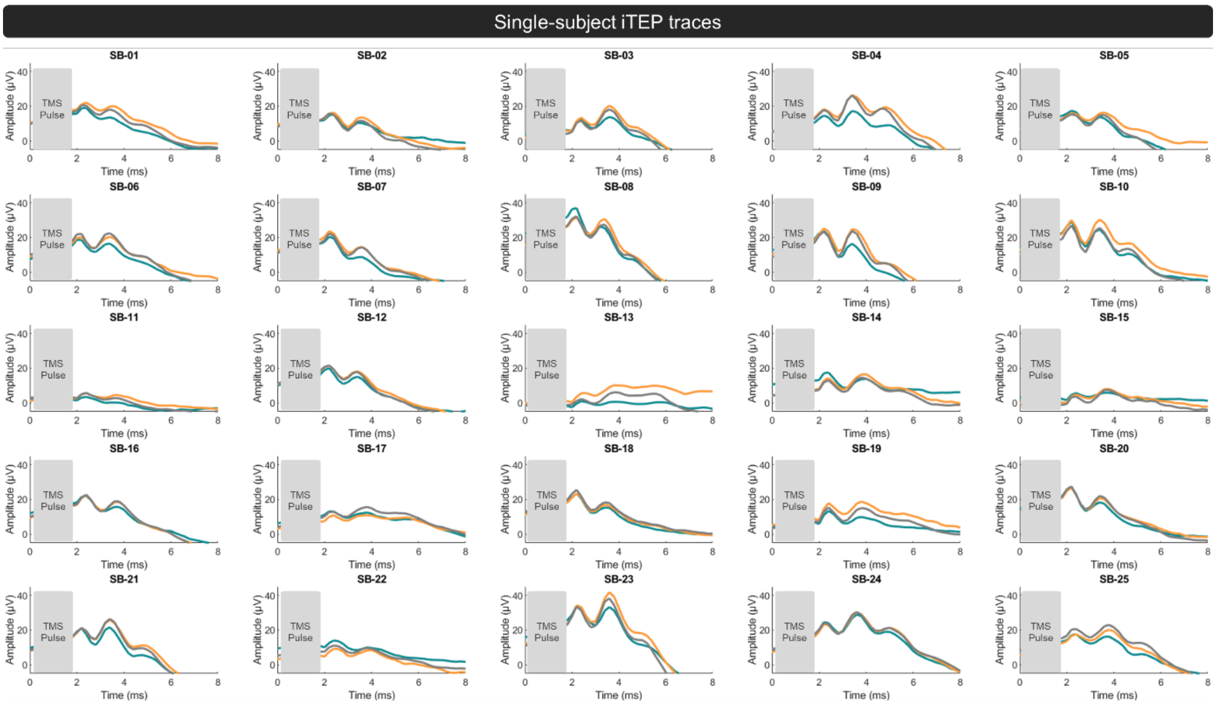


**Figure SM1.1.** Effect of movement-related beta stages on the immediate transcranial evoked potential (iTEP) peaks, visualized for each individual subject. Beta stages include movement-related beta desynchronization (MRβD; blue), post-movement beta rebound (PMβR; orange), and rest (grey). The electrode plotted is the one closest to the site of stimulation.

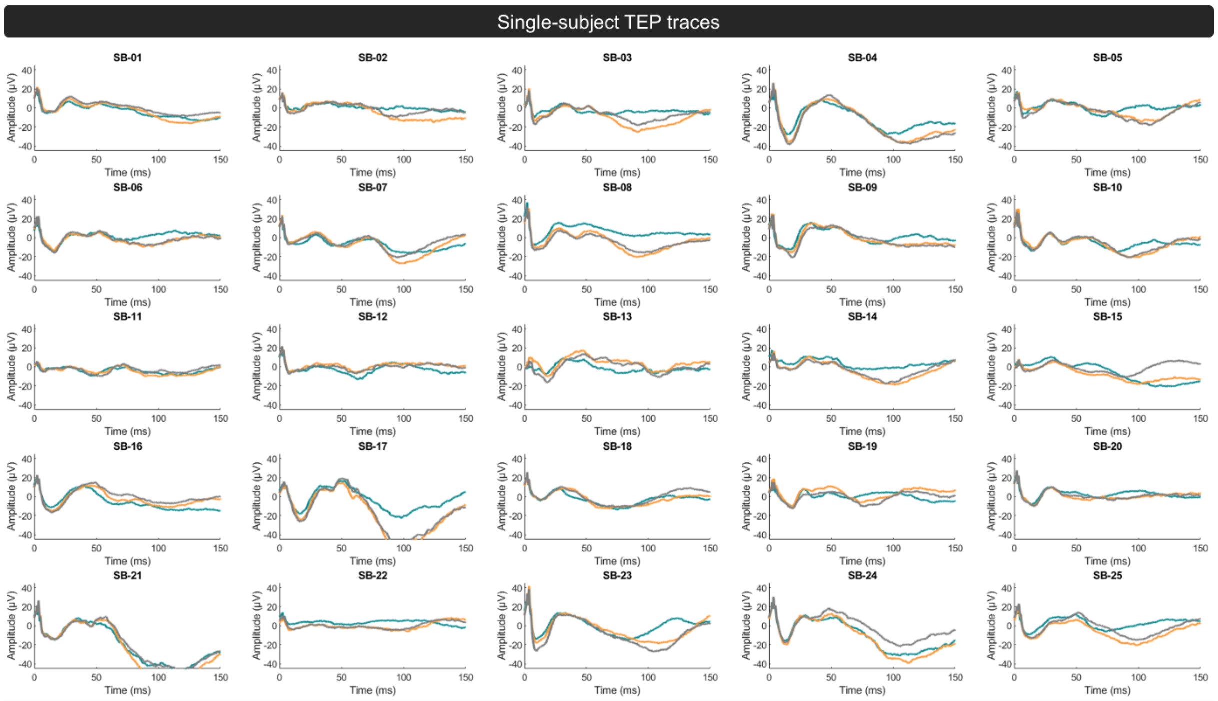


**Figure SM1.2.** Effect of intrinsic movement-related beta stages on all transcranial evoked potential (TEP) peaks, visualized for each individual subject. Beta stages include movement-related beta desynchronization (MRβD; blue), post-movement beta rebound (PMβR; orange), and rest (grey). The electrode plotted is the one closest to the site of stimulation.

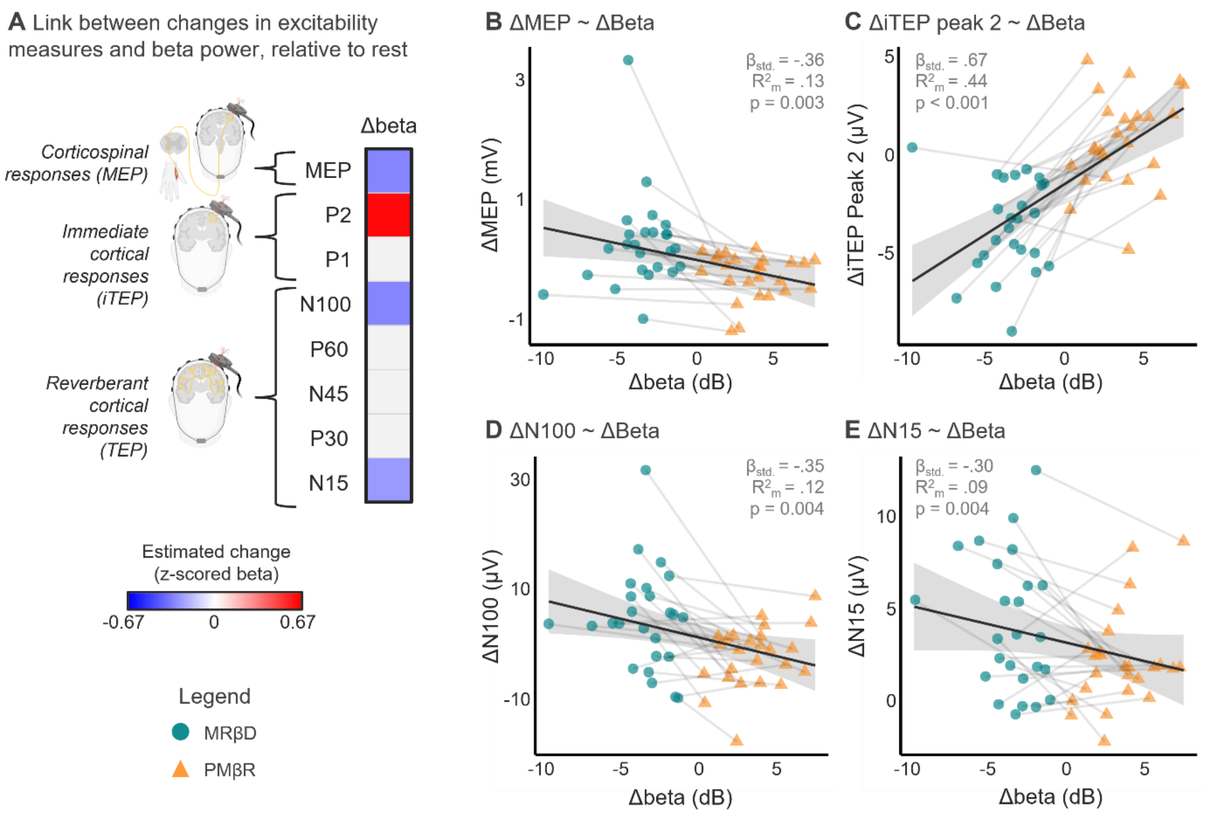


**Figure SM1.3.** Results of linear mixed effects models assessing the relationship between beta modulation with respect to rest and cortical excitability measures with respect to Rest, during two canonical beta stages; movement-related beta desynchronization (MRβD; blue) and post-movement beta rebound (PMβR; orange). Panel A provides a summary of the results, where we found significant associations between beta modulation and MEP, iTEP peak 2, TEP N100 and TEP N15. Panels B – E visualize all significant linear mixed models. A more positive ΔBeta was related to a greater iTEP peak 2 amplitude, relative to rest. Conversely, ΔMEP, ΔN100 and ΔN15 were all negatively related to ΔBeta, indicating that more beta power relative to rest, related to a smaller MEP amplitude, and larger (i.e., more negative) N15 and N100 amplitudes. Overall, these linear mixed effects models concur with the analyses presented throughout Experiment 1.

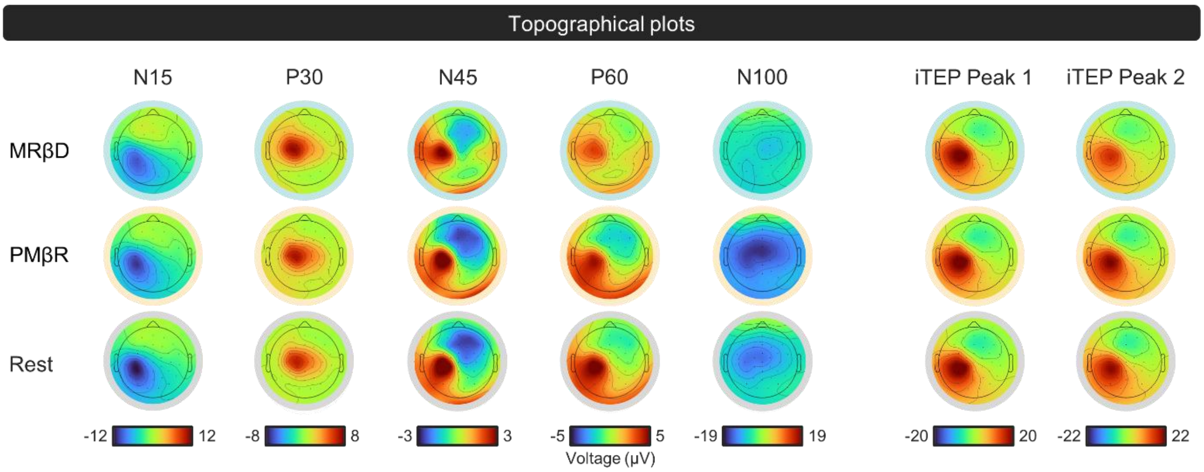


**Figure SM1.4.** Topographical distribution of mean activity at each immediate (iTEP) and later transcranial evoked potential (TEP) peak across participants, for each movement-related beta stage. Beta stages include movement-related beta desynchronization (MRβD), post-movement beta rebound (PMβR), and Rest.

**Supplementary Materials 2. Additional results and control analyses of Experiment 2.**

The second part of the Supplementary Materials provides several single-subject visualizations and additional control analyses that support the main results section of Experiment 2.

**EEG trial counts and distribution**

| Condition | Mean ± SD | Minimum | Maximum |
|---|---|---|---|
| Rest | 55.44 ± 3.97 | 49 | 60 |
| Task | 391.22 ± 25.35 | 326 | 413 |
| Catch | 64.22 ± 7.94 | 49 | 73 |

**EMG trial counts and distribution**

| Condition | Mean ± SD | Minimum | Maximum |
|---|---|---|---|
| Rest | 60 ± 0 | 60 | 60 |
| Task | 313.90 ± 37.57 | 224 | 356 |

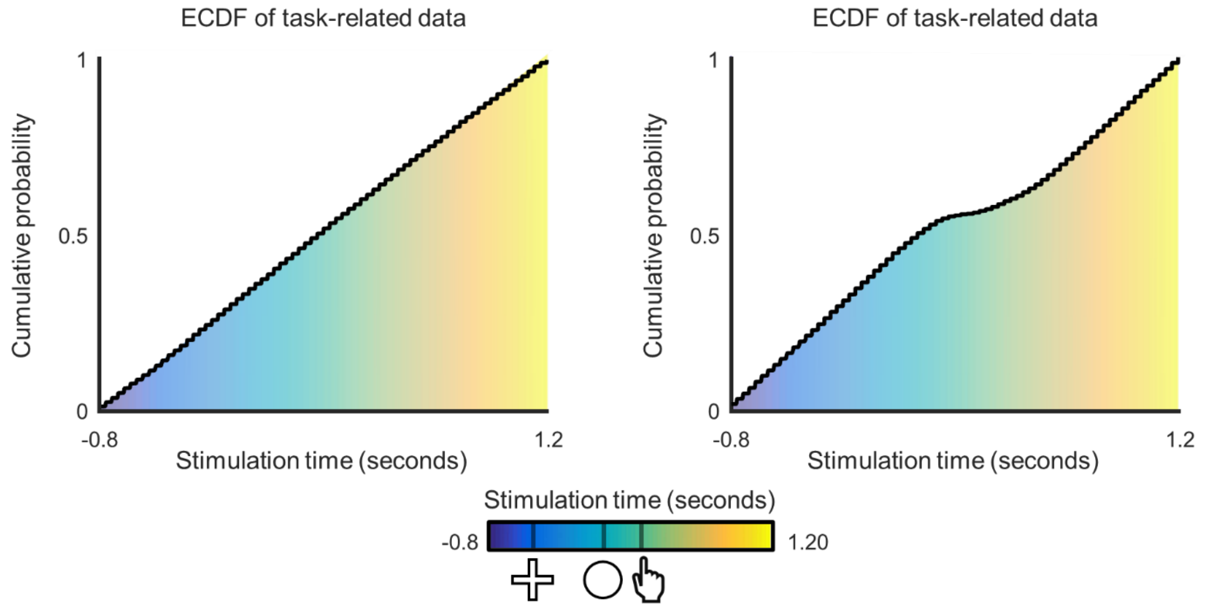


**Figure SM2.1.** The distribution of the processed electroencephalography (EEG) and electromyography (EMG) trials for Experiment 2. **Left panel**. EEG data distribution. In the upper table, descriptive statistics are provided concerning group-level trial counts for the resting-state block (pre-task), the task-related transcranial magnetic stimulation (TMS) trials with variable timings (cf., lower graph), and the catch trials (no-TMS), which were used to quantify oscillatory beta, alpha and theta characteristics and where no TMS was applied. The lower graph shows the empirical cumulative distribution function (ECDF) of the non-interpolated TMS timings with respect to the ”go” signal (0 seconds). As can be observed, the cumulative probability of the EEG trials gradually increases along with stimulation time, showcasing the success of our experimental protocol in terms of analyzing

TMS-EEG peaks across the movement cycle. **Right panel**. EMG data distribution. In the upper table, descriptive statistics are provided concerning group-level trial counts for the resting-state block (pre-task) and task-related TMS (cf., lower graph). The lower graph, the ECDF, shows that before 0 s and after 0.27 s (button press), there is a gradual increase in cumulative probability. However, between the ”go” signal and button press, there are fewer trials due to ongoing muscle activity masking the TMS pulse and subsequent MEP.

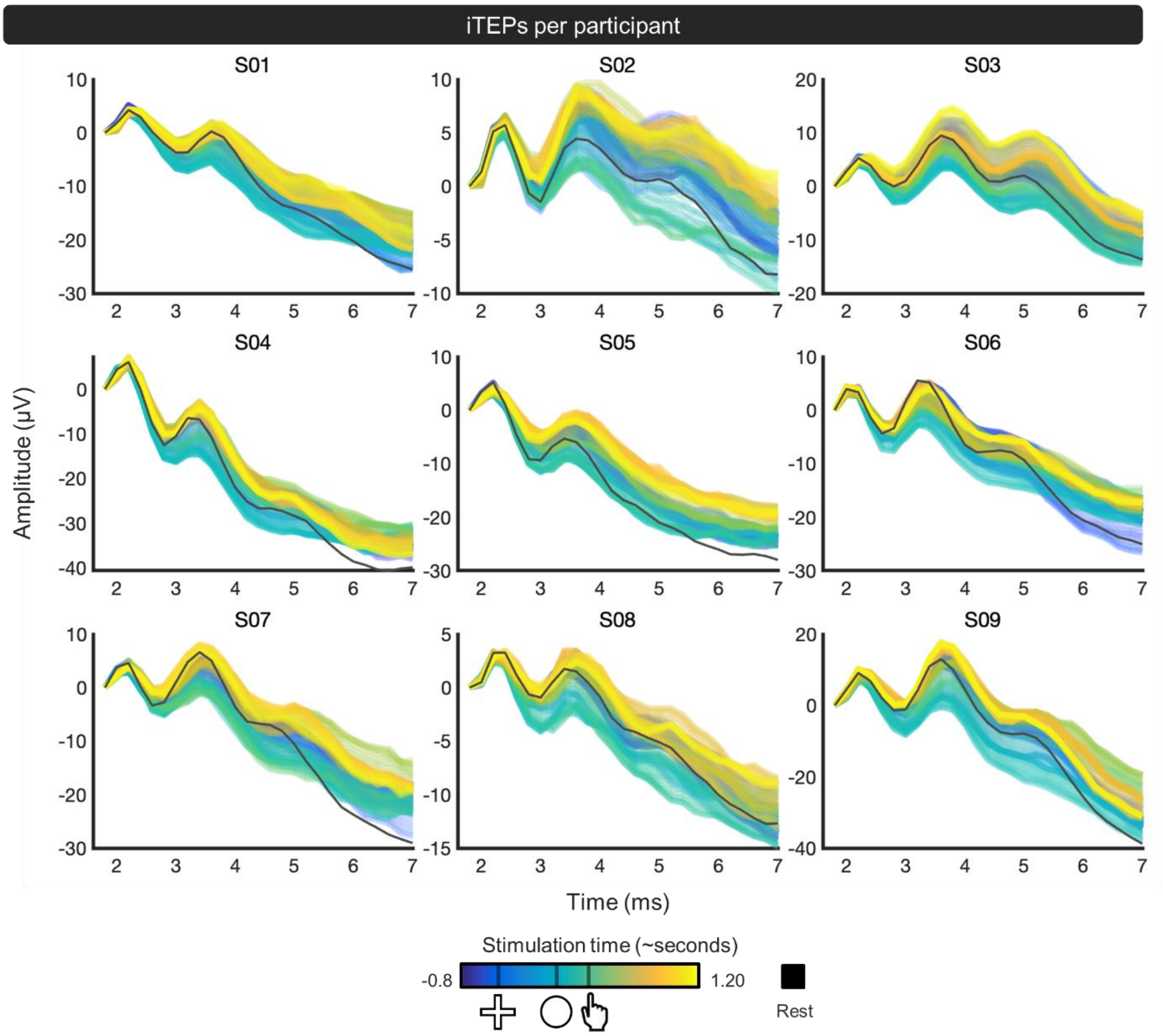


**Figure SM2.2.** Immediate transcranial evoked potential (iTEP) waveforms at rest (black) and across task-related transcranial magnetic stimulation (TMS) time points (color-coded by stimulation time), visualized for each individual subject participating in Experiment 2. The electrode plotted is the one closest to the site of stimulation. Of note, for S10, electroencephalography (EEG) data were of insufficient quality (cf., main text) and for S11, the Experiment was stopped early due to technical issues.

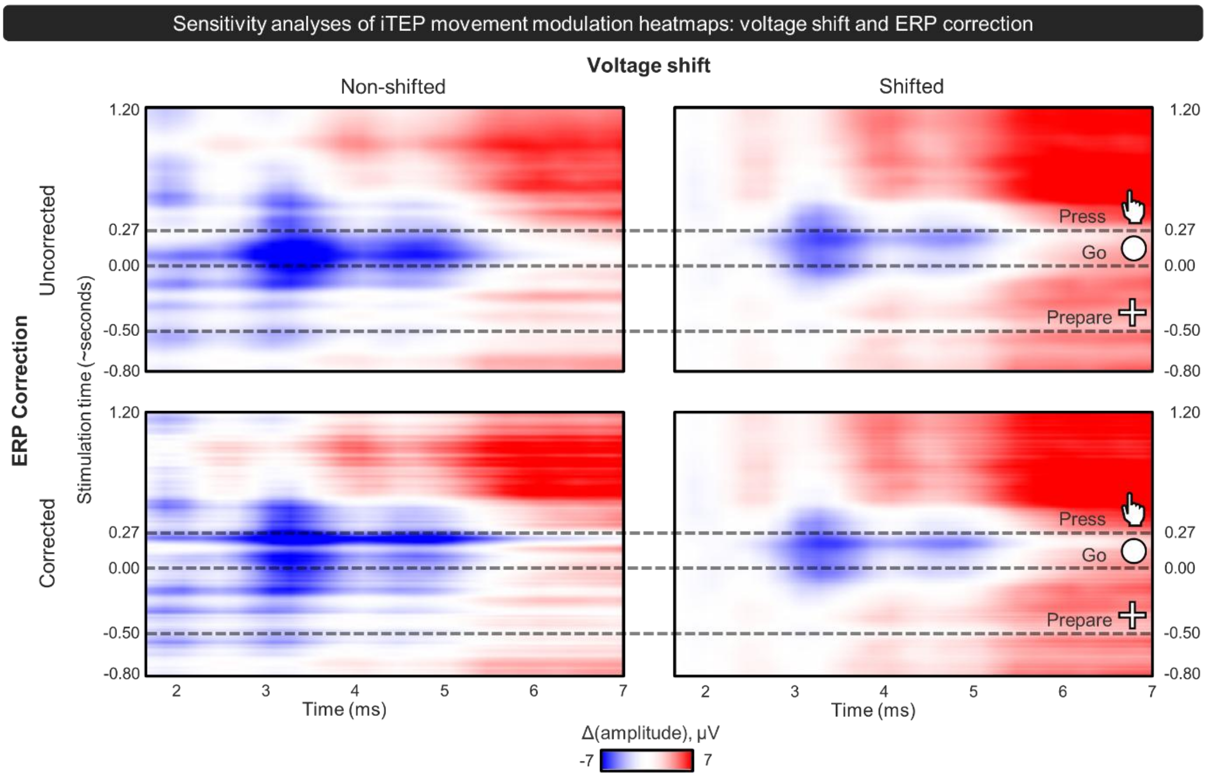


**Figure SM2.3.** Difference heatmap of immediate transcranial evoked potentials (iTEPs) relative to rest as a function of transcranial magnetic stimulation (TMS) timing (y-axis; aligned to ”go” cue) and post-TMS latency (x-axis), akin to Figure 3A. Colors indicate increases (red) and decreases (blue) in amplitude relative to rest. Dashed lines indicate task events. These heatmaps are plotted for the non-shifted (left) and shifted (right) data and without (top) and with (bottom) event-related potential (ERP) correction. The voltage shifts refers to a voltage subtraction ensuring that all iTEP traces started at a voltage of 0 μV at the first analyzed datasample (cf., Figure SM2.4.). The ERP correction refers to a voltage subtraction correcting for the ERP underlying the TMS-EEG generated iTEPs. Specifically, per participant, the ERP across the movement cycle was calculated using the same data as analyzed for the time-frequency plots in the main text (Figure 3A and 4A). The ERP, starting at the moment of TMS pulse application, was then subtracted from the iTEP data of that participant. Overall, these control analyses provide evidence that the observed movement-cycle dependent iTEP modulation are a genuine neurophysiological phenomenon, not attributable to voltage shifts or underlying ERP dynamics. Of note, the upper right figure is included in the main text (Figure 3A).

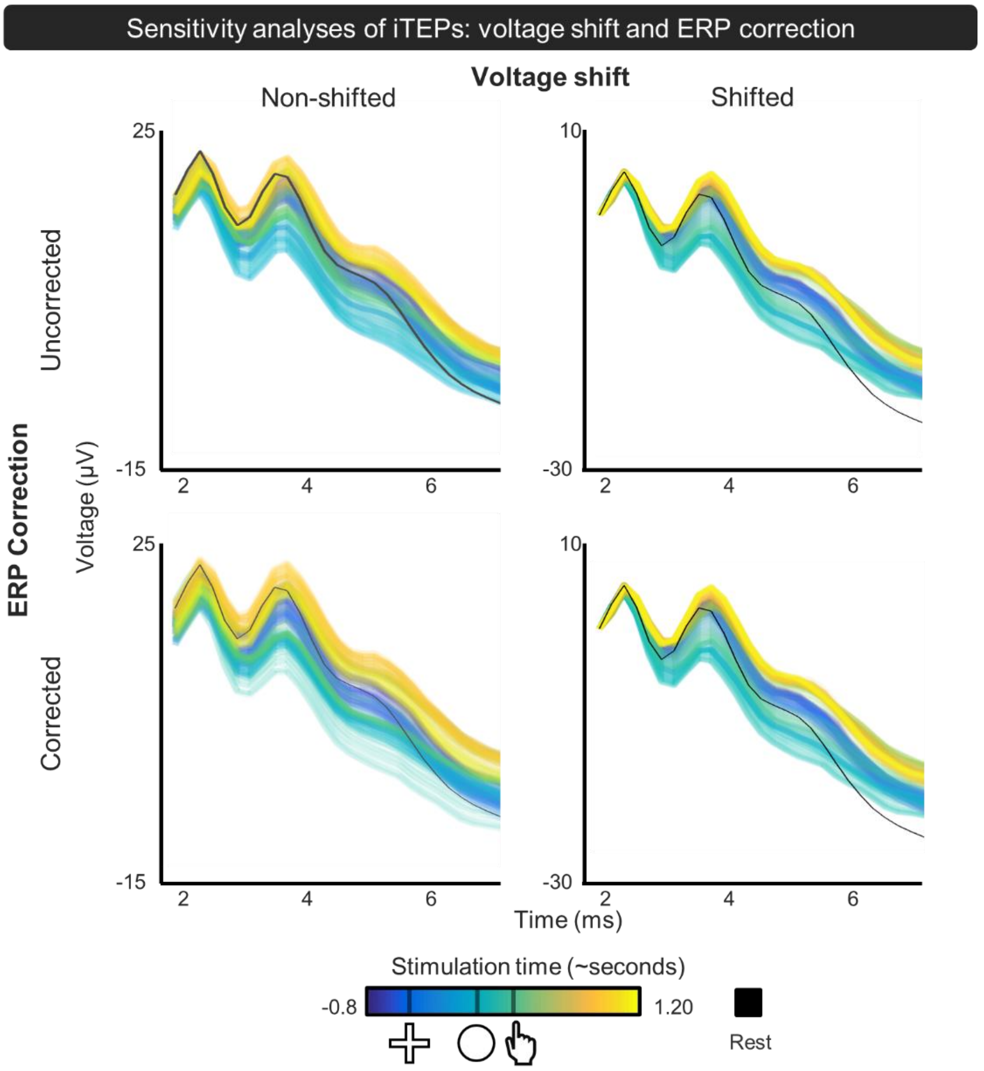


**Figure SM2.4.** Same data as plotted in Figure S2.3, now visualizing the immediate transcranial evoked potentials (iTEP) waveforms. These findings further highlight that the observed iTEP modulation is a genuine neurophysiological phenomenon not attributable to voltage shifts and/or underlying event-related potential (ERP) dynamics. Same conventions as Figure SM2.2.

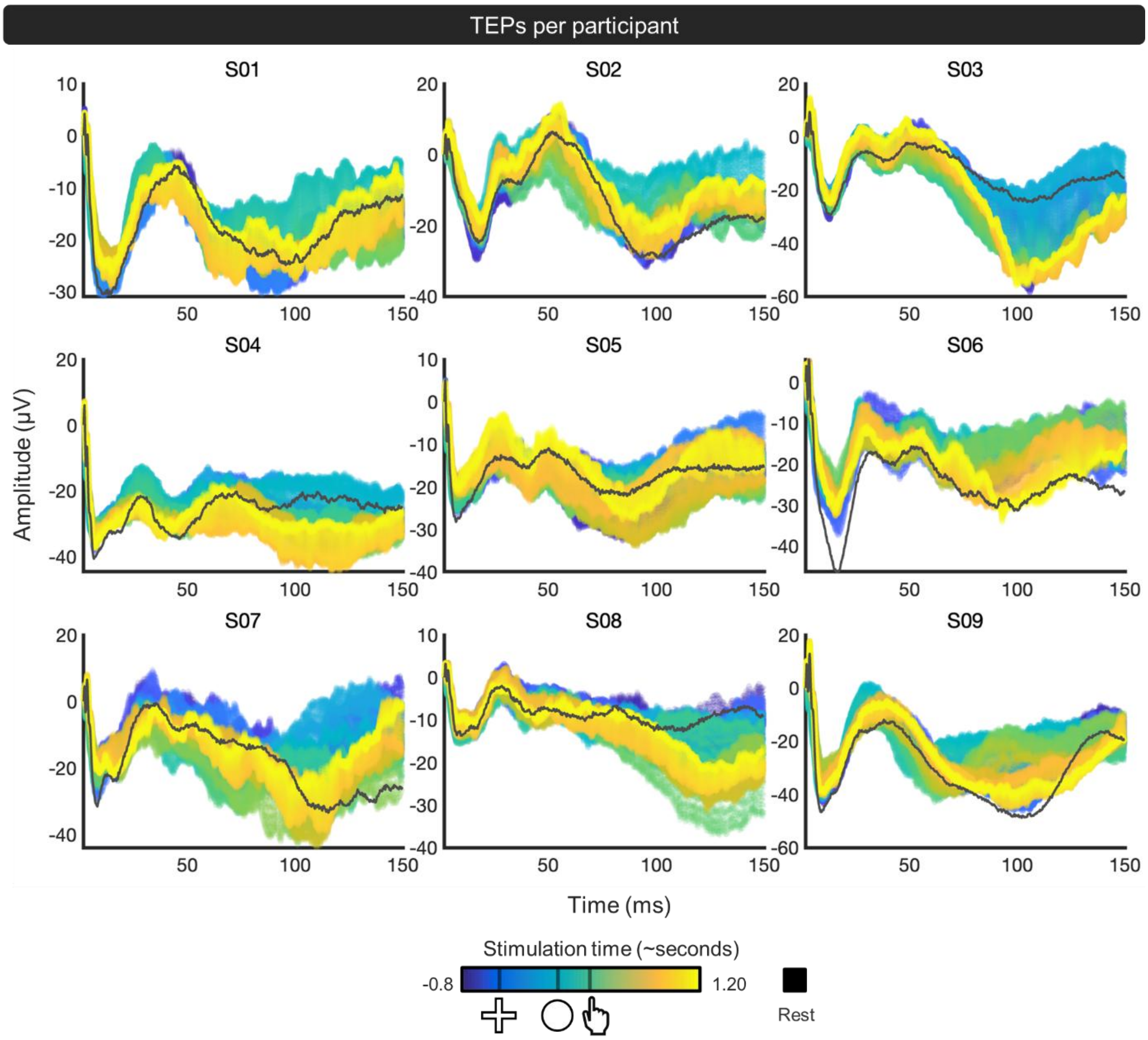


**Figure SM2.5.** Later transcranial evoked potential (TEP) waveforms at rest (black) and across task-related transcranial magnetic stimulation (TMS) time points (color-coded by stimulation time), visualized for each individual subject. The electrode plotted is the one closest to the site of stimulation. Same conventions as Figure SM2.2.

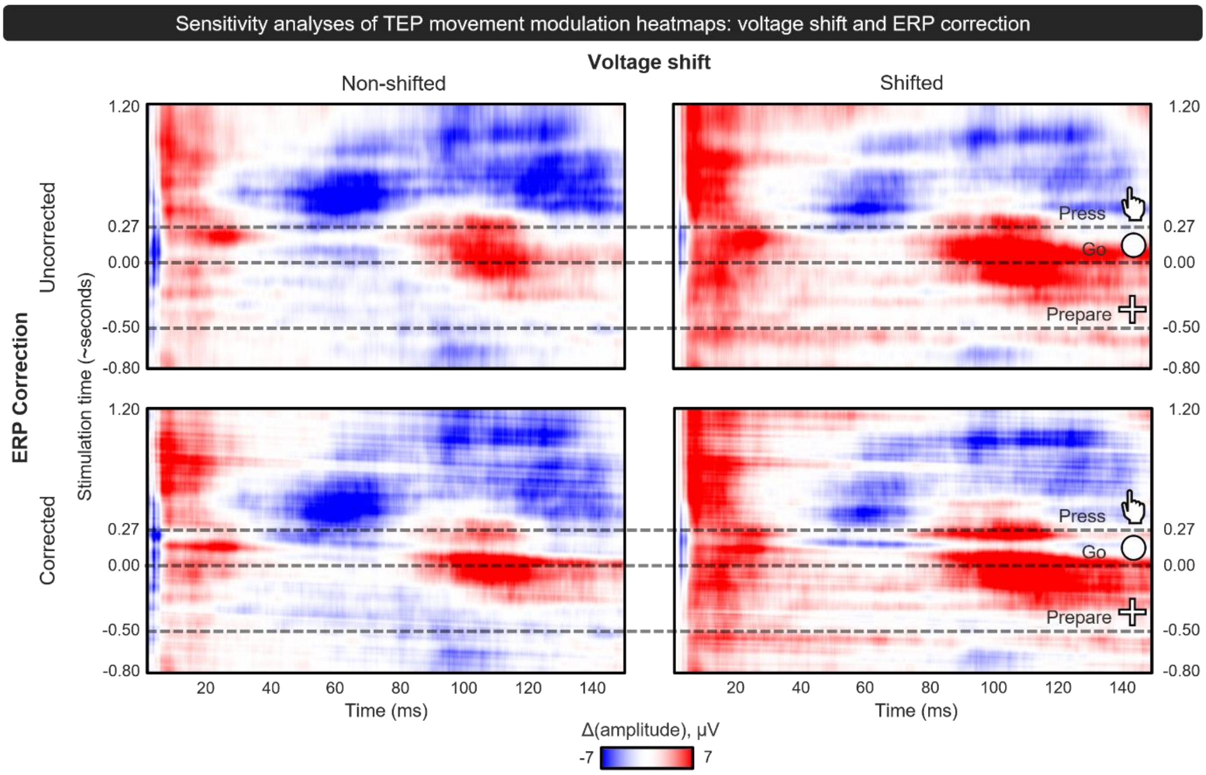


**Figure SM2.6.** Same approach as discussed in Figure SM2.3., now visualizing the transcranial evoked potentials (TEPs). Again, this highlights that the observed movement-cycle dependent TEP modulation is not attributable to either voltage shifts at the first analyzed data point or underlying event-related potential (ERP) dynamics. Of note, the upper right figure is included in the main text (Figure 4A).

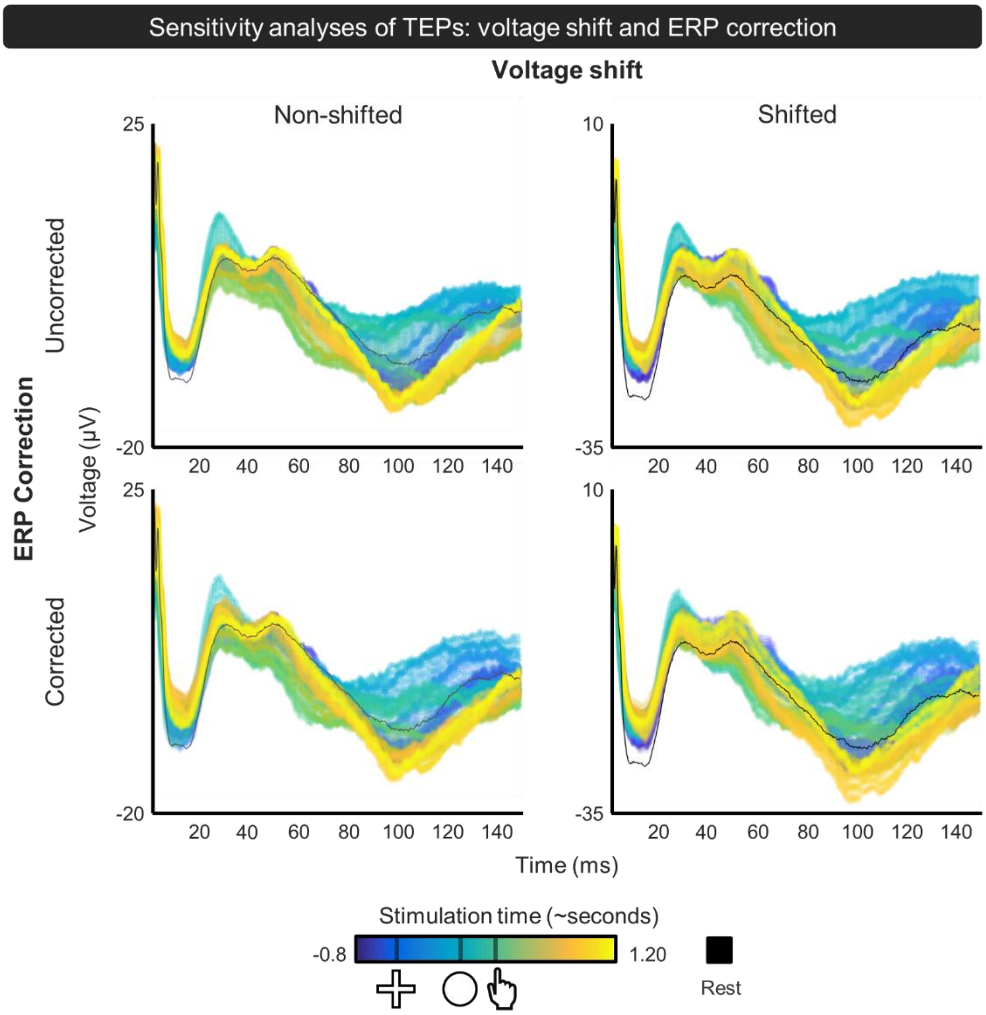


**Figure SM2.7.** Same data as plotted in Figure S2.6, now visualizing the transcranial evoked potential (TEP) waveforms. These findings further highlight that the observed TEP modulation is a genuine neurophysiological phenomenon not attributable to voltage shifts and/or underlying event-related potential (ERP) dynamics. Same conventions as Figure SM2.2.

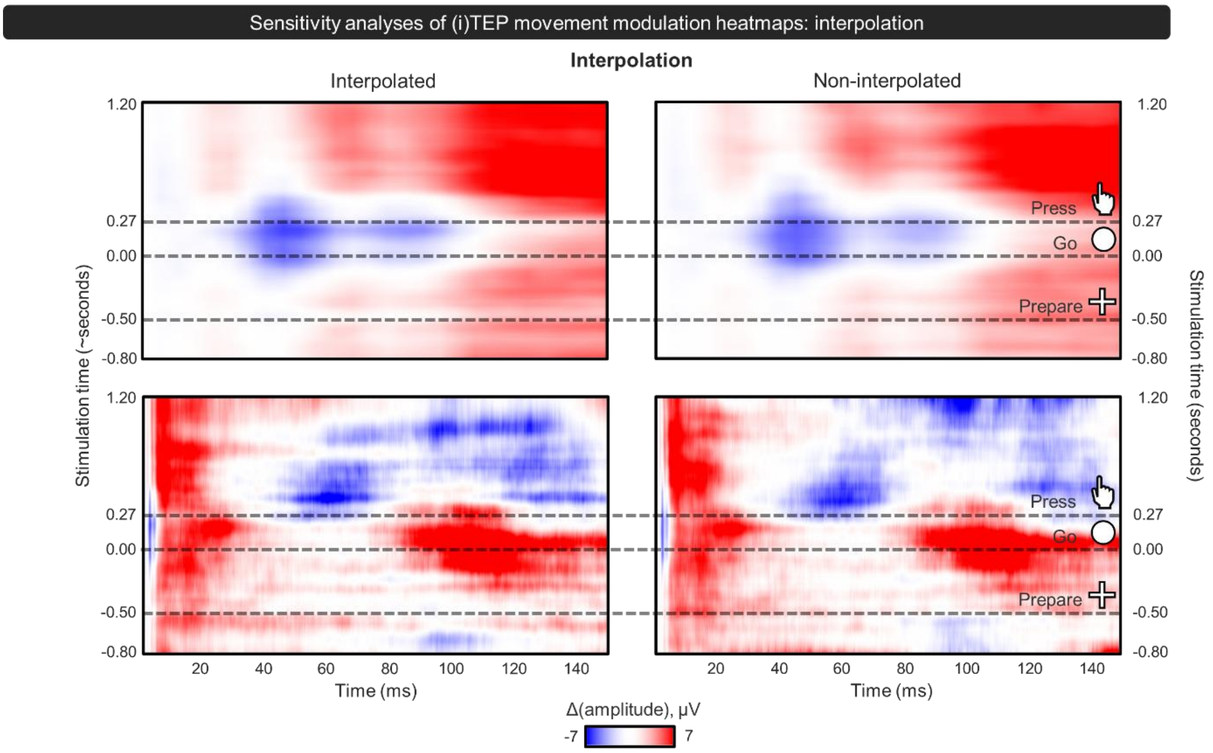


**Figure SM2.8.** Difference heatmap of transcranial evoked potentials (iTEPs, top) and TEPs (bottom) relative to rest as a function of transcranial magnetic stimulation (TMS) timing (y-axis; aligned to ”go” cue) and post-TMS latency (x-axis), akin to Figures 3A and 4A. Dashed lines indicate task events. On the left, the interpolated data is shown, as included in the main text. Specifically, the TMS-pulse timings were interpolated to ensure that the temporal intervals between trial start, go cue, and response were normalized to the group median, with the go cue fixed at 0 ms. This procedure ensures that each TMS pulse is consistently positioned relative to the underlying behavioral events (i.e., trial structure), rather than absolute time, thereby preserving its functional interpretation across trials and participants. On the right, the non-interpolated data is shown. Here, the TMS pulse was analyzed with respect to pulse adminstriation, irrespective of when this happened with respect to behavioral events. Overall, these control analyses highlight that our findings are robust to the choice of interpolation.

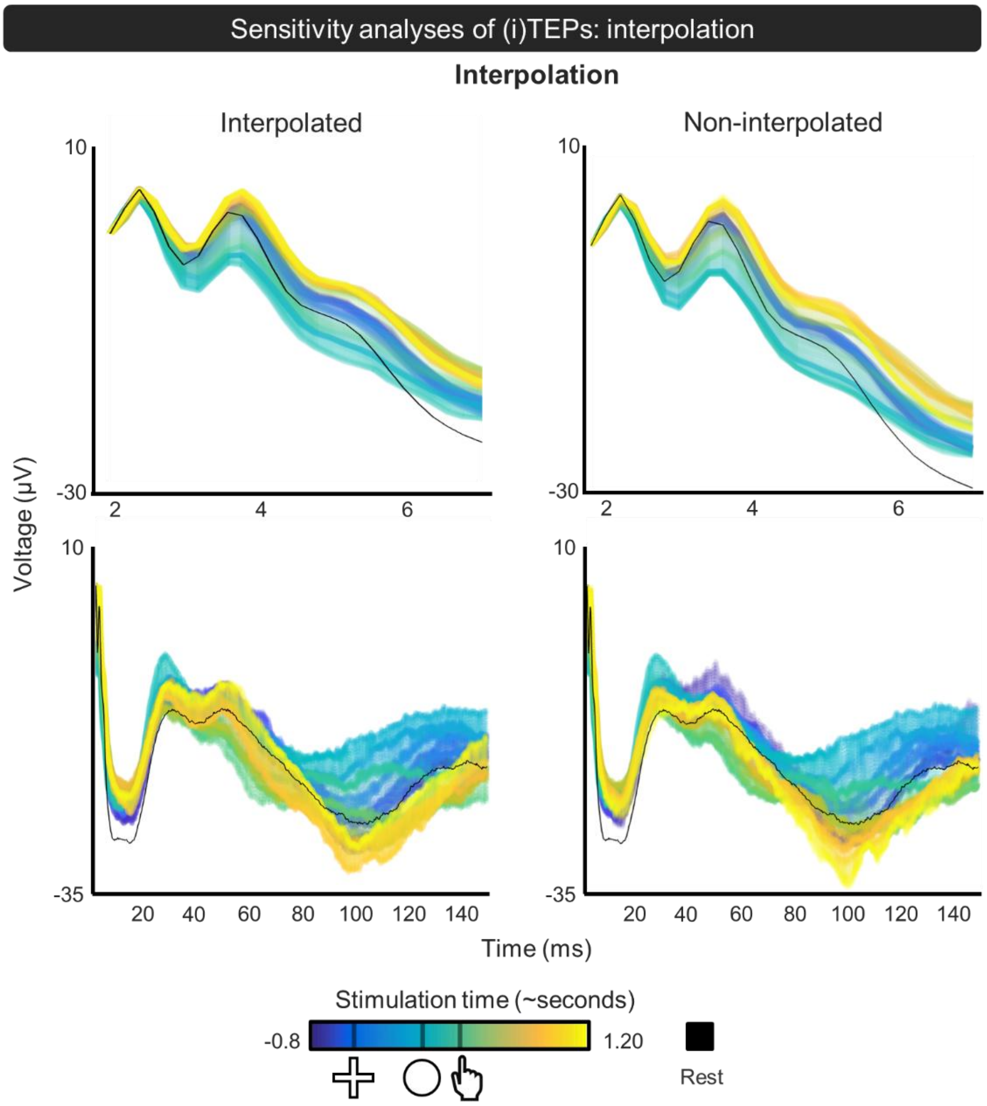


**Figure SM2.9.** Same data as plotted in Figure S2.8, now visualizing the immediate transcranial evoked potential (iTEP, top) and later TEP waveforms (bottom). Same conventions as Figure SM2.2. This further highlights that the observed (i)TEP modulation is robust to the choice of interpolation.

**Supplementary Materials 3. Targeting individual movement-related beta stages in Experiment 1.**

The third part of the Supplementary Materials provides an overview of the study's method of targeting intrinsic movement-related beta stages and extracting the individualized stimulation timings.

To personalize the timing of TMS relative to the peak movement-related beta stage, we derived participant-specific MRβD/PMβR timings from the EEG data using a standardized preprocessing and time–frequency pipeline.

Continuous EEG data were resampled to 1 kHz, band-pass filtered (1–35 Hz), average re-referenced, and denoised using ICA (Picard), with automatic component classification (ICLabel) and rejection of non-brain components (probability > 0.9). Data were epoched time-locked to the button press (–2.5 to +1.5 s). Time–frequency representations were computed per channel using complex Morlet wavelets (4–35 Hz; 31 logarithmically spaced frequencies; 3-10 cycles). Power was averaged across trials and expressed in decibels relative to a pre-movement baseline (–2.0 to –1.5 s). Analyses focused on the beta-band (13–30 Hz) within a post-cue window (50–1000 ms). Within this window, MRβD and PMβR were identified using data-driven thresholds defined by the 5th and 95th percentiles of beta-band power values, respectively. Binary masks were generated for significant MRβD (power decreases) and PMβR (power increases), constrained to the beta-band and the time window of interest. Spatially contiguous regions were labeled, and for each region the mean power change was computed. The most pronounced MRβD (most negative mean) and PMβR (most positive mean) clusters were selected. The temporal centers of these clusters (mean time index) defined the individualized MRβD and PMβR time points used to trigger beta stage-dependent TMS, which also used a 50 ms jitter.

This procedure yielded participant-specific estimates of MRβD and PMβR timing, enabling precise alignment of TMS perturbations with intrinsic motor stages. An overview of the extracted stimulation timings per subject and beta stage can be found in **Table SM3.1**.

**Table SM3.1.** Personalized TMS timings across subjects, per movement-related beta stage.

| SUBJECT | Individual TMS timings (ms) | |
|---|---|---|
| | Movement-related beta desynchronization (MRβD) | Post-movement beta rebound (PMβR) |
| SB01 | 136 | 472 |
| SB02 | 112 | 623 |
| SB03 | 93 | 766 |
| SB04 | 158 | 723 |
| SB05 | 111 | 564 |
| SB06 | 117 | 513 |
| SB07 | 129 | 714 |
| SB08 | 69 | 862 |
| SB09 | 94 | 702 |
| SB10 | 140 | 739 |
| SB11 | 132 | 800 |
| SB12 | 182 | 515 |
| SB13 | 96 | 507 |
| SB14 | 93 | 815 |
| SB15 | 80 | 721 |
| SB16 | 111 | 535 |
| SB17 | 93 | 546 |
| SB18 | 128 | 731 |
| SB19 | 75 | 611 |
| SB20 | 107 | 544 |
| SB21 | 148 | 703 |
| SB22 | 135 | 804 |
| SB23 | 113 | 542 |
| SB24 | 310 | 550 |
| SB25 | 96 | 523 |
| Median | 112 | 623 |

**Figure SM3.1** provides an overview of ongoing task-related activity across frequency bands with respect to each TMS pulse window. Several statistical checks were performed to evaluate whether our method successfully modulated and targeted beta activity. Using paired-sample t-tests, we examined differences in beta power between the baseline trials—

used to extract the stimulation timings—and the catch trials, which were intermixed with the stimulation trials. Beta values were extracted from the personalized TMS pulse window. No significant differences in beta activity were found between catch and baseline trials during MRβD ($t_{24} = -0.88$, $p = 0.39$), PMβR ($t_{24} = 1.44$, $p = 0.16$), and Rest ($t_{24} = 1.03$, $p = 0.32$) stimulation window. Beyond this, we also evaluated if beta activity in the baseline and catch windows correlated within each stage via Spearman correlations. While they correlated well in the MRβD (rho = 0.73, $p < 0.001$) and PMβR (rho = 0.59, $p = 0.002$) pulse window, no correlation was observed during Rest (rho = 0, $p = 1$). Lastly, via a mixed model, we examined if beta power during the TMS pulse window differed across the three beta stages (lmer(Beta ~ BetaStage + (1 | subject)). A significant effect of stage was found ($p < 0.0001$, $F_{2,48} = 134{,}48$, $R^2_{marginal} = 0.764$, $R^2_{conditional} = 0.790$), with MRβD (mean: -4.19 dB) being lower then Rest (mean: -0.68 dB) ($t_{48} = -8.14$, $p < 0.0001$) and PMβR (mean: 2.88 dB) ($t_{48} = -16.40$, $p < 0.0001$), and Rest being lower then PMβR ($t_{48} = -8.26$, $p < 0.0001$).

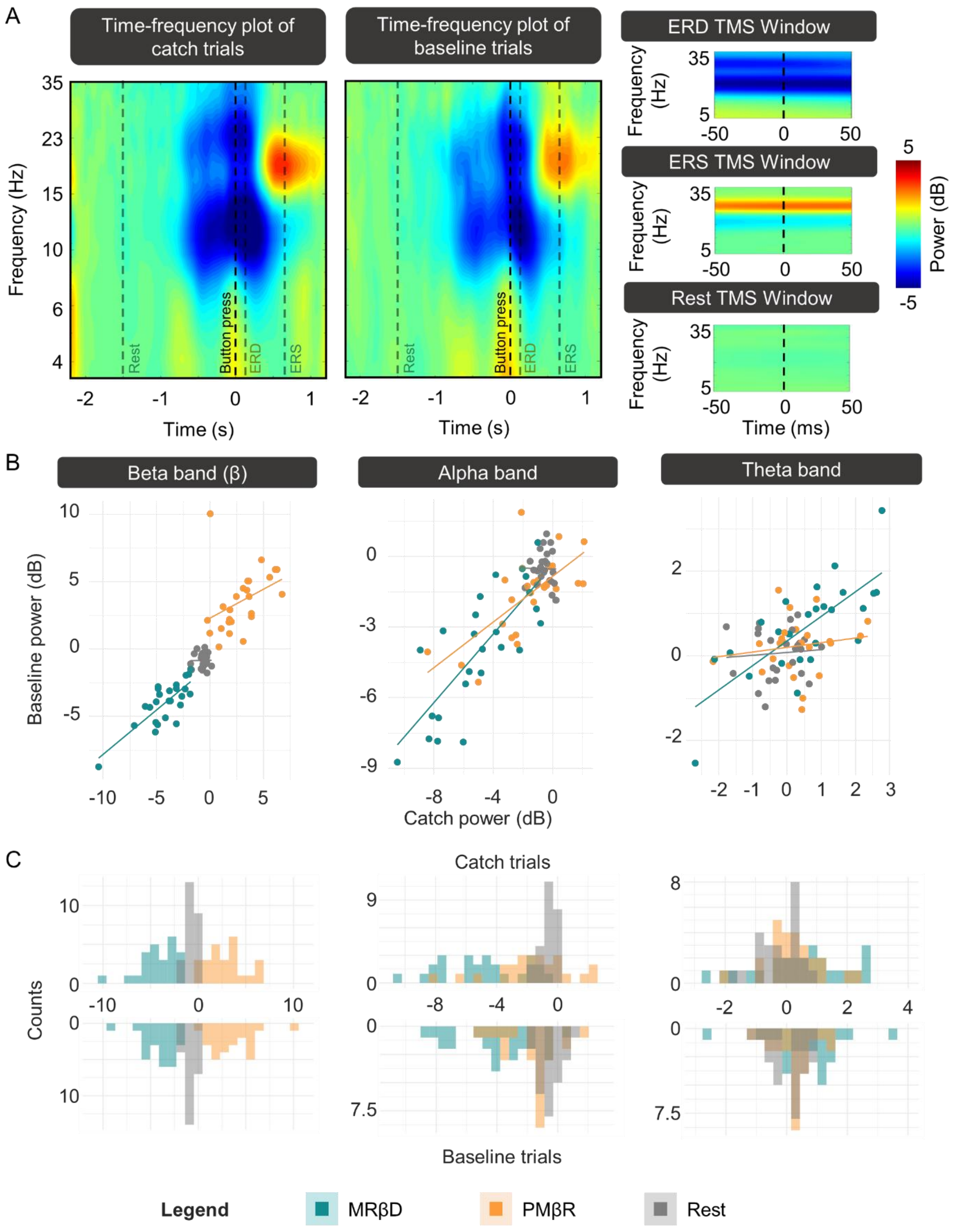


**Figure SM3.1.** Task-related modulations in beta, alpha, and theta frequency bands. **A)** Time-frequency representations from an electrode over the left motor cortex of baseline and catch trials (left), and of catch trials during the three beta stages (right). The left plots show that the reaction time task successfully modulated beta-band activity, as evidenced by clear rest, desynchronization, and synchronization stages. The right plot shows 100 ms windows, with 0 ms corresponding to the individual-specific timing of the transcranial magnetic stimulation (TMS) pulse identified from the baseline period. The -50 to +50 ms window indicates the TMS jitter. **B)** Correlation between power values in baseline and catch trials across frequency bands. Beta power showed strong correlations between baseline trials — which were used to derive personalized TMS timings— and online catch trials, indicating successful targeting. **C)** Histograms of beta, alpha, and theta power for baseline and catch trials. Power distributions were overlapping, further supporting the comparability of frequency modulations across conditions.

**Figure SM3.2** illustrates pre-stimulus EMG activity in a single trial (left) and average (right) example across the three stimulation windows. Preprocessing trial removal ensured that no residual background muscle activity was present in the 50 ms preluding TMS application.

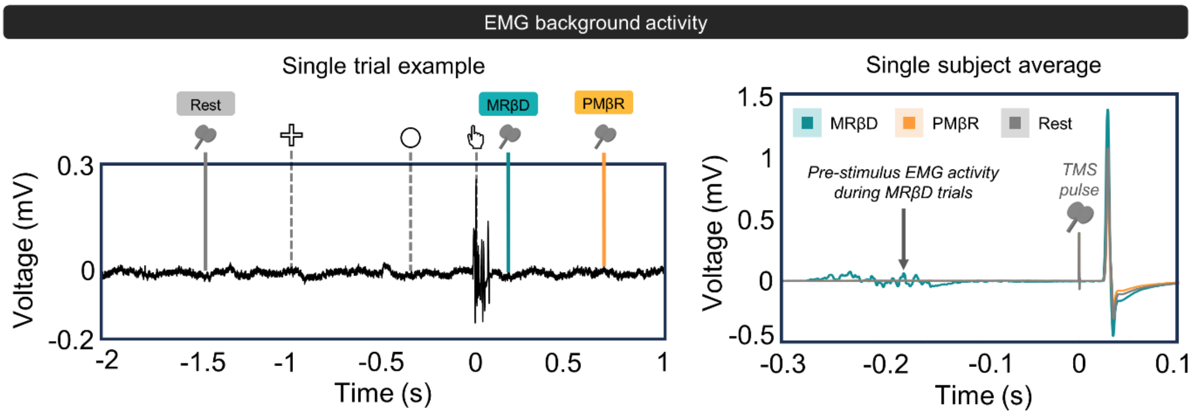


**Figure SM3.2.** Background electromyography (EMG) activity for a single representative participant. The left plot illustrates the transcranial magnetic stimulation (TMS) trigger timings relative to ongoing peripheral muscle activity in a single exemplary trial. The right plot illustrates the average (pre-stimulus) EMG activity for each beta stage.

**Supplementary Materials 4. Extracting peak TEP amplitudes**

The fourth part of the Supplementary Materials provides overview of the procedure used to extract peak and trough amplitudes for the different TEP peaks.

For iTEPs, consistent with prior work [22, 23], the amplitudes of peaks 1, 2, and 3 were identified using MATLAB's findpeaks() algorithm. For later TEP peaks, peak amplitudes were identified as absolute minima/maxima within the canonical TEP time-windows [62]. The following peaks were extracted: N15 (6–20 ms), P30 (20–35 ms), N45 (35–47 ms), P60 (47–70 ms), and N100 (70–140 ms).
